# Fixing cells with light and DNA-binding fluorophores

**DOI:** 10.64898/2026.03.27.714905

**Authors:** Maëlle Carraz, Stéphanie Bosch, Thomas Mangeat, Sylvain Cantaloube, Vincent Amarh, Romain Duval

**Affiliations:** Université de Toulouse, IRD, PHARMADEV, Toulouse, France; Université de Toulouse, CNRS, Molecular, Cellular and Developmental biology unit (MCD), Center for Integrative Biology (CBI), Toulouse, France; LITC Core Facility, Center for Integrative Biology (CBI), CNRS, Université de Toulouse, Toulouse, France; Department of Medical Biochemistry, University of Ghana Medical School, Korle-Bu, Accra, Ghana; Université Paris Cité, IRD, Inserm, MERIT, Paris, France

## Abstract

Light has become of paramount importance for the spatiotemporal control of cell labeling, activity and death. We found that palmatine, a DNA-binding alkaloid, mediates the fixation of live cells upon visible light irradiation. Fixation takes a few seconds or minutes depending on the power used, is accompanied by nuclear fluorescent labeling in target cells, exhibits a high degree of control from cell islets to single cells, yielding cells with intact nucleus and membrane boundaries. “Optofixation” relies on palmatine-DNA interaction, singlet oxygen photogeneration and lipid peroxidation, producing fixing levels of endogenous aldehydes. We also establish that many commonly used nuclear dyes exhibit optofixation properties across the UV-far red spectrum. Comprising the straightforward inactivation and labeling of native target cells by small fluorophores, this unprecedented, comparatively simple technology has the potential for major applications in biological research.

## Introduction

Chemical fixation by aldehydes, often coupled with fluorescent staining, is a ubiquitous method to arrest and stabilize biological events, with a wide range of applications in subcellular, cellular and whole organism imaging. However, exogenous aldehydes fix an entire cell population, are too slow to accurately capture fast events and generate artefacts, particularly in the nucleus.^1^ Although photoactivatable prodrugs can selectively release aldehydes in target cells, they are otherwise limited by the same drawbacks.^2,3^ Within a multicellular environment, on-demand labeling or death induction in specific cells, amongst other functions, remains pivotal to understanding cell interactions and roles in biological processes. Optogenetics, requiring prior modification of native cells,^4,5^ and laser ablation,^6^ enable precise cell targeting. Being ablative techniques, they can however modify the architecture of the systems studied. It is noteworthy that optogenetics currently lacks a fixation modality, to maintain the physical integrity of 3D microenvironments and discriminate between functional and mechanical inductions.^7^

Regarding the intrinsic biological effect of light, it depends on the wavelength, intensity, duration and temporal structure of the irradiation. Phototoxicity thus ranges from a limiting factor in live optical imaging^8^ to photosensitizer (PS)-mediated cell killing in photodynamic therapy (PDT).^9^ Rare reports of cells becoming irreversibly “frozen” or “stiffened” upon intense irradiation, yet not evolving towards complete destruction,^10-12^ were associated with light-generated reactive oxygen species (ROS) (Fig. 1B).

**Figure 1.**
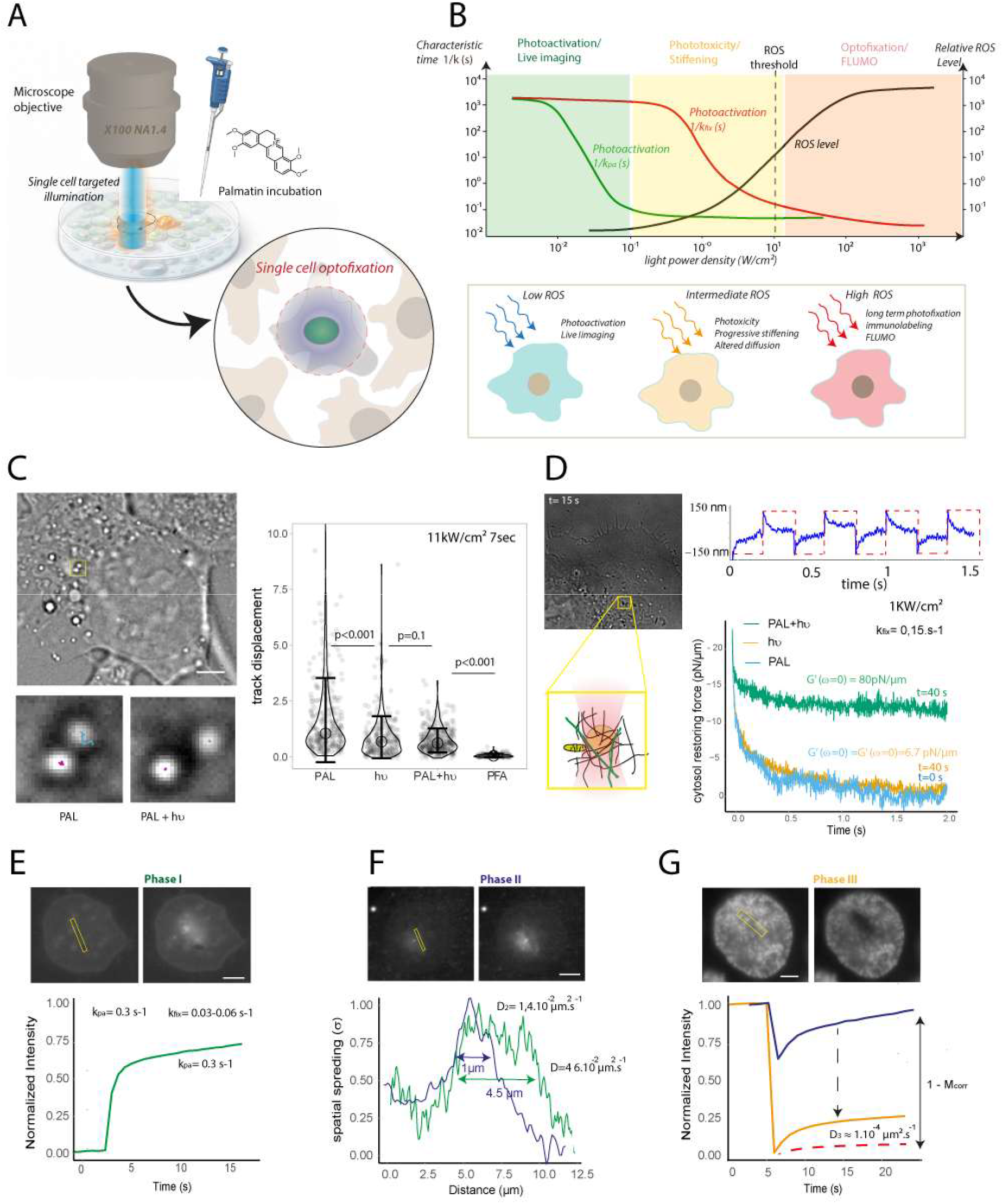
Visible-light irradiation of PAL-treated cells results in their irreversible fixation. **(A)** FLUMO principle: following an overnight incubation with palmatine, SC are fixed at 488 nm using a microscope objective. **(B)** Conceptual diagram showing the relation between time constants (1/k) and irradiation power density with cell phenotypes. **(C)** Tracking of endogenous LD by BF microscopy. *Top left*: representative cell image with tracked region of interest (yellow square). *Bottom left*: example trajectories and localization maps of vesicle mobility before (PAL) and after (PAL+hv) irradiation. *Right*: Statistical analysis of LD displacements showing a highly significant reduction in track displacement after irradiation (PAL+hv) compared to pre-irradiation (PAL), irradiation alone (hv) and FA. **(D)** OT-based microrheology showing temporal evolution of the cytoplasmic restoring force and extracted viscoelastic moduli during irradiation **(E)** Phase I: photoactivation-dominated regime. *Top*: time-lapse images showing progressive spreading of the photoactivated signal within the nucleus. **(F)** Phase II: optofixation combined with photobleaching. *Top*: the photoactivated signal displays reduced spatial spreading and altered recovery kinetics. *Bottom*: reduction of the spatial spreading from 4.5µm to 1µm. **(G)** Phase III: fully optofixed state. *Top*: time-lapse images showing slow fluorescence recovery. *Bottom*: comparison of recovery curves between 5 phase II and phase III.

We introduce here a novel general method to rapidly and selectively fix native target cells by means of photogenerated endogenous aldehydes. While screening a panel of natural fluorophores for their imaging potential in human cell lines^13^ (table S1), we found the alkaloid palmatine (PAL) to mediate an undocumented light-dependent cell fixation and labeling process. This rapid phenomenon featured complete and irreversible cell immobilization together with striking nuclear fluorogenesis (Fig. 1A), mediated by ROS, lipid peroxidation (LPO) and LPO-derived aldehydes (Fig. 1B). “Optofixation” could be conducted microscopically in various cell types under high spatiotemporal control from cell islets down to single cells (SC), yielding well-preserved, non-apoptotic fixed-labeled cells persisting for several weeks within a live-cell environment.

Beyond PAL, optofixation proved to be a property of exceptional scope, being conferrable to many DNA-binding dyes across the UV to far-red fluorescence range. Using common fluorophores and standard microscopes, fluorophore-mediated optofixation (FLUMO) constitutes a novel technological promise to quickly fix and label native target cells within complex biological systems, with spatiotemporally resolved applications in nuclear dynamics, SC, organoids and whole organism studies.

### Visible-light irradiation of PAL-treated cells results in their irreversible fixation

The FLUMO principle involves incubating palmatin in cells at a concentration of 10 µM overnight. The cells are then illuminated with a laser or LED in a targeted manner, either on cell clusters or SC (Fig. 1A), with a power range of 35 W/cm^2^ to 11 kW/cm^2^. The immediate light-induced cellular response is recorded using BF and fluorescence imaging techniques. The temperature increase associated with illumination is negligible, ranging from 0.01 to 0.3 °C (see Supplementary Materials). ***Wide-field fluorescence excitation of PAL-treated cells yields “optofixed” cells***. Unexpectedly, targeted irradiation in PAL-treated cells caused rapid and persistent arrest of intracellular dynamics, characterized by strong suppression of organelle and lipid droplet (LD) motion (Fig. 1C, left panel and mov. S1). We named this phenomenon “optofixation”. At a power density of 11 kW/cm^2^, target cells were immobilized within seconds and remained identifiable within non-target cell populations for several days following light removal and return to standard culture conditions (Fig. 1B). LD tracking confirmed this transition. Before irradiation, LD trajectories showed total displacements consistent with active intracellular transport and diffusive motion. However, after irradiation, these trajectories became strongly restricted (Fig. 1C). As previously described, irradiation alone induced only partial slowing (Fig. 1B), whereas irradiated PAL-treated cells showed near-complete LD immobilization, indicating a synergy between PAL and light exposure. ***Optical tweezer and FRAP microscopy establish the fixed nature of optofixed cells***. To confirm the increase in cytoplasmic stiffness, we conducted OT microrheology analyses on individual trapped LD in the cytoplasm. After 488nm irradiation, the elastic modulus of the cytosol increased rapidly. A 12-fold increase in the elastic component occurred over approximately 15 to 40 seconds at a power of 35 W/cm^2^ (Fig. 1D). The curves confirmed a rapid transition to a solid intracellular state under the influence of light, consistent with the observed inhibition of intracellular mobility (see Supplementary Materials). In comparison, cells fixed with formaldehyde (FA) exhibited cytosolic restoring forces that exceeded the measurement range of our OT device (>300 pN), indicating a significantly stiffer intracellular network than in optofixed cells (Fig. 1D). As a nuclear fluorogenesis with emission at 520 nm appeared to be synchronized with cell optofixation, we recorded the fluorescence after illuminating a specific region of the nucleus. The experiment continuously alternates between two stages. First, involving the diffusion of PAL following photoactivation. Second, involving optofixation induced by observation light (see Supplementary Materials). Diffusion was dominated by PAL photoactivation at the start of the irradiation (Fig. 1E). At intermediate times, the spread of fluorescence gradually slowed down, indicative of initiated optofixation. The diffusion spread was reduced 4-fold (Fig. 1F). A dominant immobile fraction in the nucleus was revealed by fluorescence recovery after photobleaching (FRAP). Indeed, diffusion of PAL was reduced by several orders of magnitude following irradiation (Fig. 1G). These results confirm a continuous transition towards a state in which chromatin is optofixed. We can also infer from these data that in optofixed cells, PAL diffuses approximately three times slower in the nucleus than LDs in the cytosol, indicating stronger fixation in the nucleus. ***Optofixed cells display normal size and morphology* and *typical DNA-mediated fluorogenesis***. The nuclear labeling of optofixed cells by PAL was attributed to its intercalation with DNA, known to be strongly fluorogenic *in vitro* ^14-16^. Despite significant inhibition of molecular motion, nuclear morphology and chromatin organization were preserved, as demonstrated by super-resolved RIM fluorescent imaging of PAL-optofixed cells *vs*. live chromatin labeling with Hoechst, DAPI or NucRed dyes^17^ (fig. S1). Collectively, our data show that PAL-mediated optofixation progresses through diffusion arrest at the subcellular scale with intracellular increase of mechanical stiffness, comprising a rapid light-driven fixation mechanism distinct from conventional chemical fixation.

### PAL-treated cells are optofixed under high spatiotemporal control, showing long-term persistence, absence of apoptotic marks and normal morphology

To better sequence and phenotypically characterize fluorogenic optofixation in cell islets, we turned to a WF microscope possessing a lower irradiation power, also capable of larger area illumination. ***PAL primarily localizes in the mitochondria but labels the nucleus upon continuous irradiation***. In accordance with previous data,^18^ PAL localizes in mitochondria prior to irradiation. However, PAL was detected as foci within MitoRed^®^-stained mitochondria ^19^, attributed to mtDNA based on the known affinity of PAL for DNA^14-16^ and similarity with PicoGreen or IF labeling of mtDNA^20,21^ (Fig. 2A). Continuous excitation of intracellular PAL (475/34 nm, 3.5 to 15 W/cm^2^ from outer limit to center of the irradiation beam) led to nuclear fluorogenic labeling in cell islets in *ca*. 4 min, proceeding *via* rapid loss of fluorescence in mitochondria then increase in the nucleus (mov. S2). This phenomenon appeared DNA-dependent^14-16^ but independent of the cell cycle (Fig. 2B & mov. S2). Our results align with the light-stimulated penetration of the archetypal protoberberine BER from mitochondria into nucleus under continuous excitation by a confocal microscope ^22^. While this phenomenon was ascribed to live-cell dynamics, we proved the light-dependent fluorogenic labeling by PAL to be entirely physicochemical as it also occurred in FA-fixed cells, albeit with basal labeling in the nucleus before photoactivation (fig. S2). This observation could be explained by a partial permeabilization of the nuclear membrane to PAL by FA,^23^ followed by an increase in permeabilization upon irradiation^11,24,25^ and ongoing optofixation altogether. ***Optofixed cells persist with a non-apoptotic phenotype***. Cells centered within the irradiation beam were strongly PAL-labeled, showing stable morphology, arrest of cell cycle and long-term persistence (Region R1, Fig. 2C & fig. S3). R1 cells were fully resistant to detergent- or cytototoxic-induced death, in contrast to surrounding live cell population (fig. S4 & S5). Despite their compromised status shown by nuclear permeabilization (fig. S6, *top panel*), R1 cells did not display early apoptosis marks (fig. S5, *bottom panel*). Optofixation could not be reproduced by sequential induction (i. e., irradiation followed by PAL treatment) (fig. S7). Taken together, our data show that optofixation and labeling are rapid events precluding apoptosis processes. Peripheral to the irradiation beam, a faintly PAL-labeled cell population was severely compromised: cells displayed shrunken nuclear morphology, permeabilization and apoptotic marks increasing overtime (Region R2, Fig. 2C & fig. S6). The delayed fluorogenesis in R2 cells was consistent with PAL nuclear uptake in dead cells (fig. S8). Surprisingly, R2 cells showed identical persistence features than R1 cells despite being severely damaged, suggesting a delayed, yet ultimately similar, endpoint fixed state (see Discussion). Concerning the relationship between optofixation and nuclear fluorogenesis, permeabilization of PAL-treated cells by Triton X-100 was fluorogenic without fixation (fig. S9), whereas fixation of PAL-treated by FA was fluorogenic (fig. S10). Consistent with dead cell staining, these results suggest that fluorogenesis is simply a marker of a fixed, permeabilized state. Accordingly, PAL accumulates in the nucleus during optofixation and fluorescence increases proportionately. ***The nuclei of optofixed cells display normal size and morphology***. Taking the cell nucleus as proxy, time-lapse imaging of PAL-mediated cell optofixing reveals a dynamic process, whereby cell nuclei undergo subtle transient expansion and contraction during irradiation (mov. S2 & fig. S11). However, SC-based size distribution analysis showed that optofixed cells from R1 display insignificant size deviation compared to the PAL-treated, live cell control (Fig. 2D). On the contrary, compromised cells from R2, as well as cells exposed to irradiation only, similarly showed severe nuclear compaction. Strikingly, FA-treated cells as positive control showed significant nuclear compaction and inferior conservation features compared to optofixed cells in R1 (Fig. 2D, fig. S11 & fig. S12). Collectively, our results show that PAL-mediated optofixation yields fully fixed, non-apoptotic, structurally maintained, fluorescently labeled cell islets, showing superior fixing quality to FA with respect to conservation of nuclear morphology.

**Figure 2.**
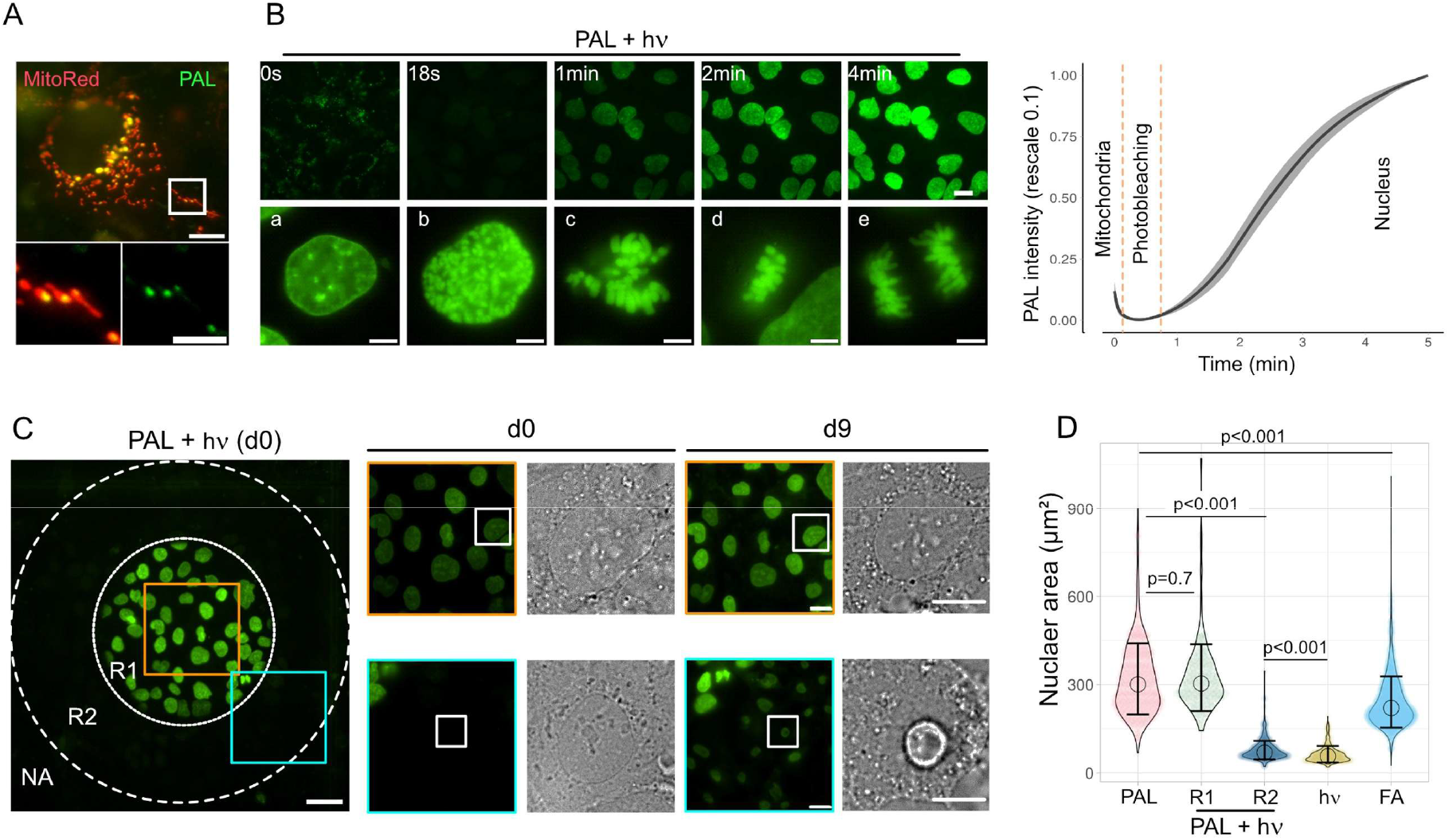
PAL-treated cells are optofixed under high spatiotemporal control, showing long-term persistence, absence of apoptotic marks and normal morphology. **(A)** Localization of PAL (green) in mtDNA within MitoRed^®^-stained mitochondria (red) in live Hep3B cells prior to irradiation (scale bar: 10 µm, crop: 5 µm). **(B)** *Top:* snapshots of time-lapse PAL nuclear fluorescence upon WF irradiation (scale bar: 20 µm). *Bottom:* PAL labeling upon irradiation at distinct phases of the cell cycle (**a**: interphase; **b**: prophase; **c**: prometaphase; **d**: metaphase; **e**: anaphase. Scale bar: 5 µm). *Right*: kinetics of PAL fluorogenesis during 5 m irradiation. Mean intensities are rescaled from 0 to 1 within a 95% confidence interval. **(C)** PAL nuclear fluorescence (in green) of a targeted cell islet following WF irradiation (PAL + hv) (R1: dotted line; R2: dashed line; NA: non-affected area. Scale bar: 50 µm). Crops in R1 (orange squares) showing stable nuclear morphology and persistence up to 9 days post-irradiation. Crops in R2 (blue squares) showing compromised nuclei (scale bar: 20 µm). **(D)** Quantification of nuclear size in different conditions: live PAL-treated cells without irradiation (PAL), FA-fixed cells (FA), PAL-free irradiated cells (hv) or PAL-treated irradiated cells (PAL + hv, R1 and R2) (n>200 per condition). Data shown in violin plots reflecting size distribution, circles indicate the medians and whiskers indicate standard deviations.

### Cell optofixation is mediated by ROS photogeneration and LPO-derived aldehydes

PAL affinity for cellular DNA (Fig. 1 & 2 and fig. S2, S3, S8, S10 & S11)^14-16^ aligns with the photophysics of protoberberines as PDT agents, known to elicit singlet oxygen (^1^O_2_) generation (type II mechanism) only as long-lived alkaloid-DNA excimers. These undergo facile intersystem crossing to the triplet state allowing energy transfer with molecular oxygen, whereas free alkaloids do not.^26-28 1^O_2_ production is associated with direct LPO^29,30^, known to yield various reactive aldehydes, mainly 4-hydroxynonenal (4HNE), malondialdehyde (MDA), acrolein (ACR), glyoxal (GA) and methylglyoxal (MGA) as breakdown products from ω-3 and ω-6 polyunsaturated fatty acids (PUFA).^31,32^ We hypothesized that LPO-derived aldehydes could be the endogenous fixing effectors in optofixed cells *via* covalent bond formation with proteins.^31^ ***Optofixation correlates with aldehyde generation and protein cross-linking***. We first investigated the biochemistry of a whole optofixed cell population using a portable LED lamp to mimic microscope irradiation (fig. S13). 467 nm LED exposure of PAL-treated cells yielded a homogenous population of fluorescently labeled fixed cells within 15 min (fig. S14). These conditions were compared to the action of representative aldehydes at fixing concentrations. We used 4% FA (*ca*. 1.33 M) as a positive control and found 1 mM 4HNE or ACR capable of fixing cells, in contrast to MDA, GA and MGA. Following cell lysis, severely reduced protein contents were measured in cells treated with FA, 4HNE, ACR or PAL/LED, major effects being observed for 4HNE and ACR (table S2). These were associated with loss of detectable soluble proteins, aberrant high molecular weight bands in SDS-PAGE, together with important insoluble loads consistent with cross-linked protein adducts^31^ In particular, the electrophoretic profile of PAL/LED cells was similar to that of 4% FA. In the absence of PAL, LED-irradiated cells showed normal protein content and SDS-PAGE profile (Fig. 3A). Second, we tested specific scavengers of LPO-derived aldehydes to inhibit PAL-mediated fluorogenesis and cell fixation. Fluorogenesis kinetics and nuclear size quantification revealed that optofixation was strongly inhibited by pyridoxamine (POA) and carnosine (COS) (Fig. 3B & fig. S15), consistent with aldehyde trapping by these amines.^31,33^ ***Optofixation correlates with strong LPO***. We assessed the LPO status of optofixed cells using the BODIPY 581/591-C_11_ redox probe.^34,35^ Irradiated PAL-treated cells underwent strong LPO in both R1 and R2, the oxidation ratio increasing up to 9-fold in R1 30 min post-irradiation relative to non-irradiated cells (Fig. 3C & fig. S16). Irradiation in the absence of PAL also led to LPO, estimated to contribute to 20-25% LPO in R1 (Fig. 3C). Increased oxidation in R1 was transient and decreased post-irradiation (fig. S13). Generic antioxidants and specific LPO inhibitors were incapable of inhibiting fluorogenic optofixation even at high concentration (see Supplementary Materials), apart from the vitamin E analogue lazaroid U-83836E (LAZ), inducing delayed fluorogenesis and impaired nucleus size preservation (Fig. 3B & fig. S15). Our results are consistent with an LPO-dependent optofixing phenomenon occurring at a rate such that it can only be buffered, but not prevented, by high concentrations of lipophilic antioxidants. ***Optofixation correlates with specific ROS production***. We tested normoxia (5% O_2_) as well as cysteamine (CEA), a specific quencher of fluorophore triplet state used in super-resolution imaging,^36,37^ to verify that optofixation-associated LPO is a molecular oxygen-dependent phenomenon. While lowering free oxygen was ineffective in preventing optofixation, cell fixation and fluorogenesis were strongly inhibited by CEA (Fig. 3B & fig. S15). CEA-pretreated cells showed extinguished nuclear labeling post-irradiation, evolving overtime to a severely compacted nuclear phenotype. Interestingly, the latter was reminiscent of PAL-treated cells having undergone insufficient irradiation for optimal optofixation (R2, Fig. 2C & 2D). Lastly, we tested the ROS Brite^TM^ 670 fluorogenic probe, possessing a 30-fold selectivity for the superoxide anion (O_2_°^-^) and hydroxyl radical (OH°) compared to ^1^O_2_ ^38^ Irradiated PAL-treated cells underwent ROS increase only up to 2-fold in R1 relatively to non-irradiated cells, while irradiation in the absence of PAL led to similar levels as in R1 (Fig. 3D). These data indicate that O_2_°^-^ and OH°, albeit generated during optofixation, arose primarily due to irradiation alone in the absence of PAL. Collectively, our results suggest that optofixation is a PAL- and irradiation-dependent phenomenon resulting mainly from ^1^O_2_ production and to a lesser extent from light-induced O_2_°^-^ and/or OH°, together responsible for strong LPO and *in cellula* generation of lipid-derived aldehydes.

**Figure 3.**
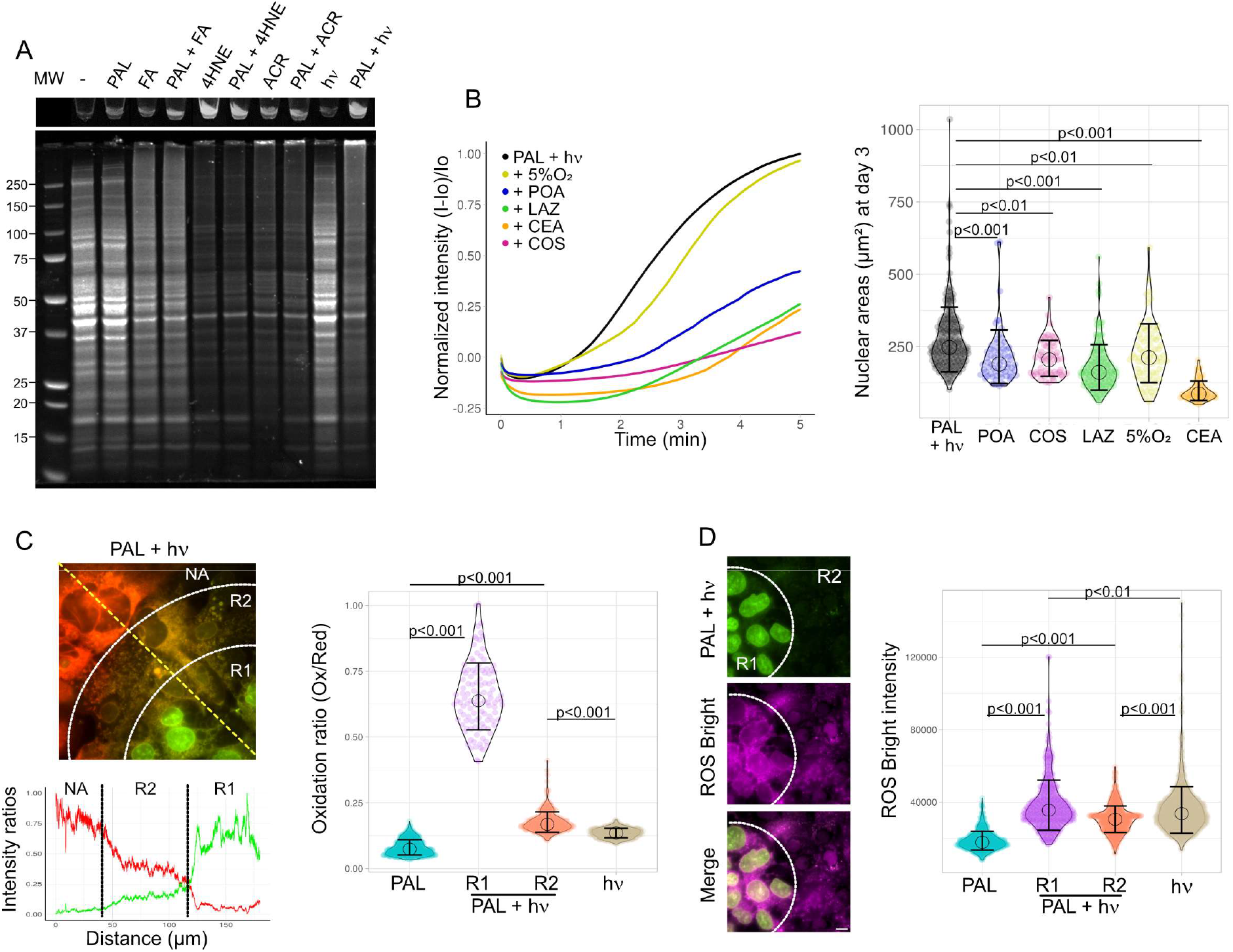
Cell optofixation is mediated by ROS photogeneration and LPO-derived aldehydes. **(A)** Biochemical analysis of a whole cell population optofixed by a LED lamp. *Top:* Visualization of abundance of precipitates (UVA irradiation). *Bottom:* SDS-PAGE analysis of soluble protein contents following treatment with FA, 4HNE, ACR or LED irradiation in the presence or absence of PAL. **(B)** Influence of optofixation inhibitors. *Left:* fluorogenesis kinetics during 5 m irradiation in the presence of various inhibitors. *Right*: quantification of nuclear sizes in the same conditions (n>60 per condition) 3 days post-irradiation. (**C)** Lipid peroxidation (LPO)-quantification in PAL-treated cells using BODIPY™ 581/591-C11 probe. *Top left:* representative image of probe staining 30 m after irradiation of PAL-treated cells in zones NA, R1 and R2. *Bottom left:* plot of ratiometric measurements along the yellow dashed line. Red curve represents Red/Ox ratio, green curve represents the Ox/Red ratio. *Right:* quantification of the oxidation ratio (Ox/Red) 30 min after irradiation of PAL-treated cells (R1, R2) compared to irradiated cells in the absence of PAL (hv) or cells treated with PAL without irradiation (PAL) (n>100 per condition). **(D)** ROS quantification in PAL-treated cells using ROS Brite^TM^ 670 probe. *Left:* representative image of ROS staining (in magenta) obtained after irradiation of PAL-treated cells (in green) in R1 and R2 (scale bar: 10 µm). *Right:* Quantification of ROS Brite fluorescence intensity.

### Optofixation constitutes a novel tool in cell biology

We investigated the biological and chemical scope of optofixation and its compatibility with state-of-the art methodologies in cell biology. ***PAL-mediated fluorogenic optofixing is observed in various cell types***. Optofixation could be achieved by PAL in cultures of diverse cell types, either normal or cancerous, mammalian or non-mammalian, suggesting the robustness and scope of FLUMO in a wide range of biological systems (Fig. 4A). ***Optofixation allows native IF***. Based on their fixed status, the feasibility of native (i. e., FA- and detergent-free) IF labeling in optofixed cells was investigated, to demonstrate a possible assessment of cell types and protein expression status in addition to their functional ablation. Representative nuclear markers of euchromatin (H3Ac), heterochromatin (SUZ12), interspace chromatin (PML bodies), telomeres (TRF1) and nuclear membrane (lamin B1) were fully detectable following native IF labeling in R1 cells (Fig. 4B). These yielded similar patterns to standard FA fixing whereas non-irradiated cells remained unlabeled, establishing the compatibility of optofixation with native IF analysis in the nucleus (Fig. 4B). As an example, native IF labeling of lamin B1 in optofixed cells showed very high similarity with RIM imaging of the protein, corresponding to its well-known pattern. However, certain other nuclear markers showed less homogeneous labelling in R1 cells (fig. S17A), while cytoskeletal proteins displayed partially expected patterns, suggesting epitope lability during optofixing (fig. S17B). To gain further insight into IF labeling in optofixed cells, the photostability of representative proteins was studied by Western blot analysis of LED-irradiated whole cell populations. Antigens showed variable photostability depending on irradiation time, irrespective of their cell localization (fig. S18). Taken together, these observations suggest that native IF in optofixed cells is possible but prone to case-by-case optimization, i.e. by modulating irradiation power and time. ***Optofixation is achievable at the SC level***. Fixing a SC selectively without affecting neighboring cells within a whole population constitutes a major new challenge in cell biology. Using a WF microscope equipped with a TIRF imaging module, we were able to irradiate SC at high power (11 kW/cm^2^) without interfering with surrounding cells (Fig. 4C). To achieve this, we set the ASTER illumination system to the minimum surface area and irradiated a SC for 8 s targeting its nucleus, yielding strongly PAL-labeled SC whilst neighboring cells showed no fluorescent labeling. The videos produced (mov. S1) illustrate the power of this methodology, the speed of fixation and the precision of irradiation limited to the SC scale. ***Optofixation is exclusively attributable to DNA-binding fluorophores across the UV-far red spectrum***. To expand the scope of FLUMO, we investigated other fluorescent trackers featuring diverse spectral features and target organelles. Using their optimal excitation wavelengths, optofixing was achievable as expected with higher-energy, i.e. blue-shifted, but also lower-energy, i.e. red-shifted fluorophores relative to PAL. However, these fluorophores were strictly limited to cytopermeable DNA dyes. Thus, other natural protoberberines e.g. BER and JAT, and benzo-δ-carboline alkaloids e.g. CRY and NMC, together with archetypal nuclear trackers e.g. DAPI and Hoechst 33342, were identified as powerful optofixing reagents and could be categorized as either fluorogenic or non-fluorogenic (Fig. 4D & fig. S19). As observed with PAL, optofixed cells exhibited preserved nuclear morphology in R1 (Fig. 4D & fig. S19), whereas R2 cells showed severe nuclear compaction or lack of labeling (fig. S20). Regarding dyes known to be genuinely nucleus-impermeable in live cells, e.g. protoberberines, since fluorophores need to be DNA-bound to elicit ^1^O_2_ photogeneration, it must be assumed that optofixation was triggered by an initial photopermeabilization of the nuclear membrane ^11,24,25^ followed by a further increase in permeabilization under both irradiation and ongoing fixing.^23^ Non-cytopermeable DNA dyes (e. g. PI, 7-AAD, SYTOX^TM^ Blue) were incapable of optofixing cells (table S3). Interestingly, cells transiently expressing GFP or mCherry-tagged proteins in the nucleus were unable of self-optofixing (fig. S21), consistent with the reliance on DNA-associated PS to induce optofixation over nuclear localization of the fluorophore alone. In full contrast to DNA-binding dyes, trackers of other cell compartments completely lacked optofixing capabilities (table S3).

**Figure 4.**
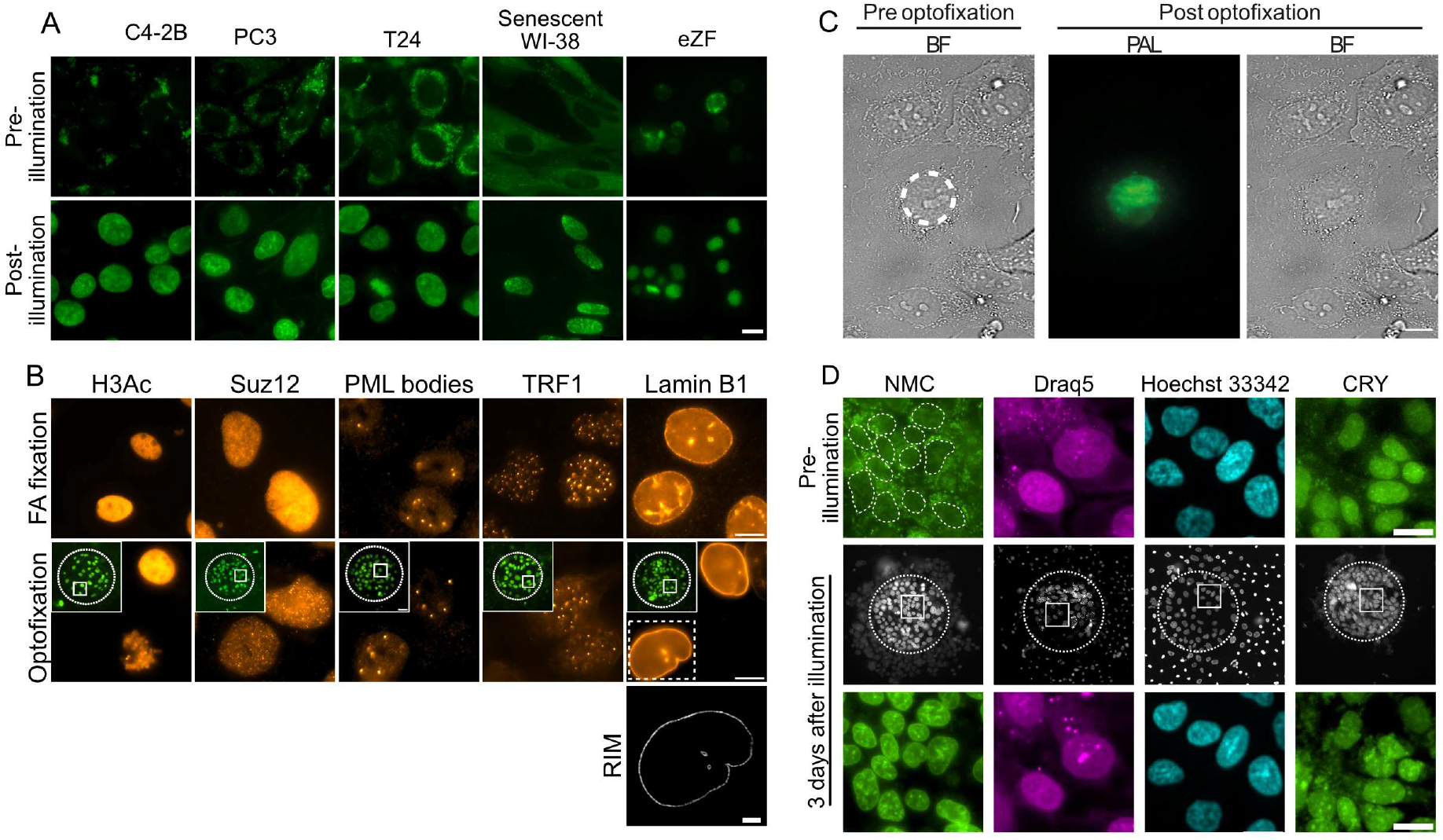
Optofixation constitutes a novel tool in cell biology. **(A)** PAL-mediated fluorogenic optofixation in human proliferative cancer cells, human senescent cells and embryonic zebrafish dissociated cells (eZF, 48 hpf) (scale bar: 10 µm). **(B)** Optofixation is compatible with native IF labeling in PAL-treated cells. Cell imaging shows PAL staining after optofixation (green) and targeted antigens (hot orange). Inserts show round dotted line (R1 cells). For each tested antigen, native IF post-optofixation (*bottom*) is compared to standard FA-based IF (*top*) (scale bar: 10µm). For lamin B1, a super-resolved RIM image after optofixation and native IF illustrates the conserved pattern. **(C)** Optofixation of SC. Dashed circle indicates the irradiated SC. The green and orange hot images show that PAL labelling is prominent in the nucleus of the targeted cell, whereas surrounding cells are unlabeled. (scale bar: 10 µm). **(D)** Examples of nuclear trackers with optofixing properties. Crops in R1 cells (dashed circle) are observed before and 3 days after irradiation (scale bar: 10 µm).

FLUMO represents a new general phenomenon of light- and DNA-dependent, fluorophore-mediated fixation of live cells. Using microscope irradiation, it allowed for the targeted fixing and fluorescent labeling of cells under precise spatiotemporal control down to SC. Revealed using the alkaloid PAL, optofixation is a common property of various natural or synthetic DNA-binding dyes across the UV-far red fluorescence range, paving the way for the functional repurposing of many classical fluorophores. While irradiation in the presence of PS supposedly leads to cell death such as in PDT, we found that high-intensity irradiation resulted in rapid cell fixation, kinetically surpassing cell damage and preserving morphological features such as cell boundaries and nucleus. With respect to the destruction of mitochondria under the conditions used (mov. S1 & S2), the fate of other cytosolic compartments during optofixing remains to be investigated in detail. We wish to emphasize that optofixation clearly is a non-linear phenomenon, influenced by physical (power and time relationships), photochemical (fluorophores and their DNA affinity, as well as triplet state properties) and biological (cell types and their PUFA composition) features. Variations in these parameters and their effects on the photophysical response of biological systems remain to be fully studied.

From a photobiological standpoint, fluorophore-treated but insufficiently irradiated R2 (non-target) cells, as well as untreated cells undergoing full irradiation, were fully compromised with alterations in global morphology including nuclear compaction. However, R2 cells persisted for days in culture with a similar fixed state as for R1 (target) cells. These observations suggest that effective exogenous PS e.g. PAL, under low irradiation, or weak endogenous PS, such as many fluorescent co-factors e.g. flavines, under strong irradiation, do generate ROS and LPO-derived aldehydes but in a manner incompatible with optimal cell fixation.^10,11^ Optofixation-associated LPO in R1 was mainly PAL-dependent and ^1^O_2_-induced through PAL-DNA complexes,^26-28^ accompanied by a minor contribution (20-25%) of light-induced ROS (mainly O_2_°^-^ and/or OH°) *via* putative cell fluorophores. Considering a combined presence of endogenous fluorophores and an exogenous DNA dye, our proposed model for FLUMO rests on the properties of excited PS in aerobic conditions, being (i) the abstraction of hydrogen radicals from biomolecules including lipids (type I mechanism, M1) leading to biomolecule oxidation and radical ROS and (ii) the production of ^1^O_2_ (type II mechanism, M2).^9,29,30^ LPO resulting in a common output, we hypothesize that the efficiency of live-cell FLUMO relies on the conjunction of both pathways (Fig. 5).

**Figure 5.**
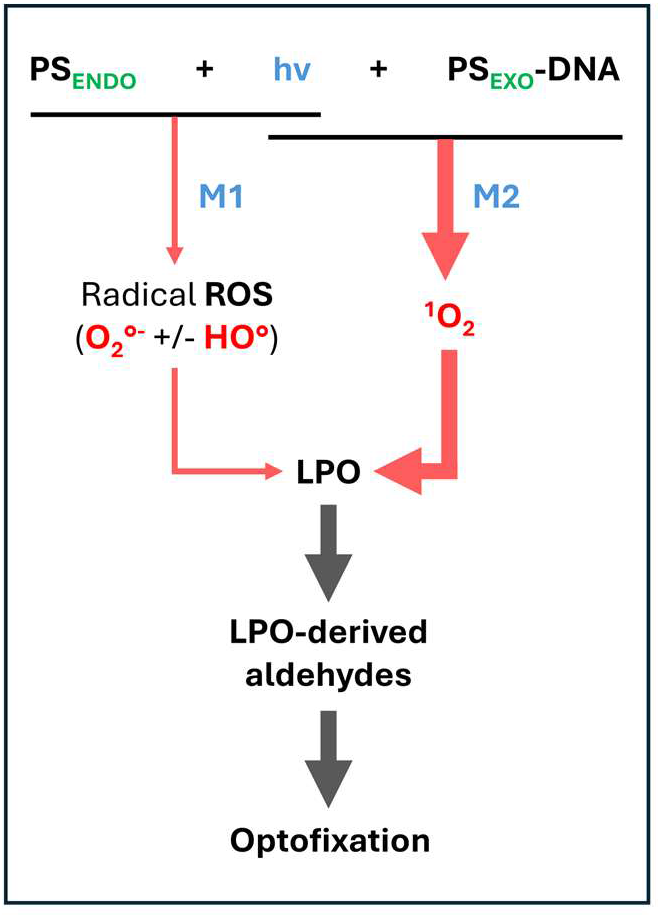
Proposed model for live-cell FLUMO

Which entities could constitute the PUFA fuel and propagate optofixing inside a living cell? ^1^O_2_ can only diffuse within a 20 nm radius^39^ to yield neutral lipid hydroperoxides (LH) by direct addition to PUFA.^29,30^ Consequently, chromatin-associated lipids,^40^ as well as nuclear and mitochondrial membrane lipids in direct proximity to PAL-DNA complexes, would be primarily subject to peroxidation. LH recently emerged as diffusible LPO “embers” *via* the endoplasmic reticulum (ER) and then Golgi apparatus to the whole cell in ferroptosis.^41^ A plausible scenario for optofixation could thus comprise in the diffusion of LH from the nucleus and mitochondria into the ER *via* their closely connected membrane systems.^42,43^ Minor in quantity but photogenerated in a pan-cellular manner, O_2_°^-^ and/or OH° would locally propagate LPO by radical cascades, yielding lipid aldehydes as end-products.^44-46^ Accordingly, the diffusion of LH and their local breakdown into aldehydes, rather than aldehydes being produced in a few major sites i.e. nucleus and mitochondria then diffusing, is likely to account for whole-cell FLUMO both quantitatively and kinetically. Mitochondria would likely contribute very little in the initiation, due to their lower DNA content limiting the rate of ^1^O_2_ generation,^26,27^ but also considering their rapid destruction during optofixing, preventing local photocatalysis. Diffusion experiments (Fig. 1C, D, E, F & G and supplementary text) and microscopic observations (mov. S1 & fig. S11) do support that optofixation is driven from nucleus to cytosol but require further evidence.

Technologically, there is no known equivalent to FLUMO, its generality regarding nuclear dyes as well as operational simplicity setting it apart from photoactivatable aldehyde prodrugs^2,3^ or optogenetically controlled cell death.^5^ Its scope in molecular life science thus appears promising across three main areas. First, to capture elusive nuclear events and their dynamic structural information. Second, in SC biology for the selective fixing and labeling of either SC or irrelevant surrounding cells, greatly facilitating downstream analyses. Third, in organoid and developmental biology to decipher cell interaction and roles: while the maintenance of 3D structures post ablation remains a major issue^6^, optofixation would consist in the functional but non-physical ablation of target cells within complex environments, a goal not yet met by genome editing. We also show that optofixation could be reproduced by an external LED source on whole fluorophore-treated cell populations. The repurposing of DNA fluorophores as easy-to-use and safe optofixatives, capable of fixing and labeling cells in a short single step, consists in a valuable alternative to classical procedures. Our results should also call for great caution regarding fixation biases when imaging live-cell or macromolecule dynamics in the presence of nuclear trackers, especially when high irradiation powers are used.

## ABBREVIATIONS

7AAD: 7-aminoactinomycin
ACR: acrolein
ASTER: adaptative scanning for tunable excitation rendering
BER: berberine
BODIPY: boron dipyrromethene
BF: bright field
CEA: cysteamine
CPG: CytoPainter^™^ Green
CRY: cryptolepine
DAPI: 4’,6’-diamidino-2-phenylindole
EpCAM: epithelial cellular adhesion molecule
ER: endoplasmic reticulum
FA: formaldehyde
FBS: fetal bovine serum
FLUMO: fluorophore-mediated optofixation
FRAP: fluorescence recovery after photobleaching
GA: glyoxal
GAPDH: glyceraldehyde-3-phosphate dehydrogenase
H3-Ac: pan-acetylated histone H3
H3K9Me3: *N*-trimethylated lysine residue 9 in histone H3
H3K27Ac: *N*-acetylated lysine residue 27 in histone H3
4HNE: 4-hydroxynonenal
hpf: hours post-fertilization
IF: imunofluorescence
JAT: jatrorrhizine
LD: lipid droplet
LED: light-emitting diode
LPO: lipid peroxidation
MDA: malondialdehyde
MEM: minimum essential medium
MGA: methylglyoxal
NA: non-activated
NMC: *N*-methyle cryptolepine
NucRed: NucRed^™^ Live 647
NucView: NucView^®^ 530 Red Caspase-3 Dye
PAL: palmatine
PDT: photodynamic therapy
PI: propidium iodide
PML: promyelocytic leukemia
PUFA: polyunsaturated fatty acid
R: region
RIM: random illumination microscopy
ROS: reactive oxygen species
SC: single cell
SDS-PAGE: sodium dodecylsulfate polyacrylamide gel electrophoresis
SUZ12: polycomb repressive complex 2 subunit
TIRF: total internal reflection fluorescence
TRF1: telomeric repeat binding factor 1
WF: wide field.

## ACKNOWLEDGEMENTS

Prof. Patrick Kobina Arthur (Department of Biochemistry, Cell and Molecular Biology, University of Ghana, Accra, Ghana) is acknowledged for facilitating project initiation between Ghana and France (NKABOM mobility fellowship to VA, 2021). RD is grateful to Dr. Yaw Aniweh, Prof. Theresa Manful and Prof. Gordon Awandare (West African Centre for Cell Biology of Infectious Pathogens, WACCBIP, University of Ghana, Accra, Ghana), as well as Dr. Jean-François Franetich (Centre d’Immunologie des Maladies Infectieuses, Sorbonne Université, Paris, France), for participating to the screening of natural fluorophores. The authors are thankful to Prof Kerstin Bystricky and her team (CBI, UT, Toulouse, France), for their intellectual inputs, plasmids’ courtesy, ensuring successful completion of the project and reviewing the work; to Dr Julie Batut (CBI, UT, Toulouse), for discussions and providing zebrafish material; to Christian Rouvière (CBI, UT, Toulouse), for his intellectual input on cell image analysis; to Prof. Patrick Kobina Arthur and Dr. Adrian J. F. Luty, for syntax corrections and suggestions to improve the manuscript; to Dr. Valérie Jullian (UMR 152 IRD PHARMADEV, Toulouse, France), for her kind facilitation of the study over the years. LITC imaging facility is member of Genotoul-TRI and the National Infrastructure France-BioImaging (https://ror.org/01y7vt929) supported by the French National Research Agency (ANR-24-INBS-0005 FBI BIOGEN).

## FUNDING

The Ambassade de France au Ghana (Accra, Ghana) is acknowledged for providing financial resources (NKABOM mobility fellowship 2021, recipient VA). UMR 261 MERIT IRD (Paris, France) is acknowledged for funding a pilot screening of natural fluorophore by cell imaging (SPACES funding, recipient RD). UMR 152 PharmaDEV (Toulouse, France, recipient MC), MCD-CBI (Toulouse, France, Tremplin funding, recipients K. Bystricky and MC) and CBI-LITC (Toulouse, France, TIRIS Strategic Booster program funding, recipient SC) are acknowledged for their large participation with supplying materials and with costs of cell imaging experiments.

## AUTHOR CONTRIBUTIONS

(ICMJE and CRediT guidelines were used to define authorship and contributions) Conceptualization: RD, MC, SB, TM

Data curation: SB, MC, TM

Formal analysis: SB, MC

Methodology: MC, SB, RD, TM

Resources: MC, SB, TM, RD, SC

Investigation: MC, SB, RD, TM, SC, VA

Visualization: SB, MC, TM, SC

Funding acquisition: RD, MC, TM, SC

Project administration: MC, RD

Supervision: RD

Writing – original draft: RD, TM

Writing – review & editing: RD, MC, SB, TM, SC, VA

## COMPETING INTERESTS

The authors have no competing interests to declare.

## DATA AND MATERIALS AVAILABILITY

All data are available in the main text or the supplementary materials.

## SUPPLEMENTARY MATERIALS

Supplementary tables S1-S3, movies S1 & S2, figures S1-S21 are available online.

## Notes

### Competing Interest Statement

The authors have declared no competing interest.

### Summary of Updates

Title updated; abstract updated, introduction updated; main results updated; discussion updated; supplementary informations updated

