## Supplementary information for "Fixing cells with light and DNA-binding fluorophores"

### Supplementary Materials

**Chemicals and reagents**.

Natural fluorophores were obtained from Merck (guaiazulene, piperine, atractylodin, anacardic acid, (15:3) anacardic acid, lasalocid sodium salt, β-hydrastine hydrochloride, kynurenic acid, xanthurenic acid, alternariol, alternariol monomethyl ether, coumestrol, (+/-)-heraclenin, oxypeucedanin, byakangelicol, byakangelicin, colladin, farnesiferol A, farnesiferol C, rutaecarpine, evodiamine, tryptanthrin, palmatine chloride, berberine chloride, jatrorrhizine chloride, quinine hydrochloride) or synthesized according to published procedures (cryptolepine triflate, *N*-methylcryptolepine iodide) [1,2]. Reference fluorescent trackers for cell imaging were obtained from Merck (DAPI, Hoechst 33258, Hoechst 33342, propidium iodide, 7-aminoactinomycin (7-AAD), Nile Red), Invitrogen (DRAQ-5, SYTOX™ Blue, ER Tracker™ Blue-White, LysoTracker™ Red, NucView® 530 Red Caspase-3 Dye, CellMask™), Abcam (CytoPainterGreen), Thermo Fisher (NucRed™ live 647, BODIPY™ 581/591 $C_{11}$) or Dojindo (MitoRed®). 4-hydroxynonenal (4HNE) was from VWR-Avantor. Malondialdehyde (MDA), glyoxal (GA), methylglyoxal (MGA), *L*-carnosine (COS), pyridoxamine (POS), (*D, L*)-α-tocopherol (TOCO) and cysteamine (CEA) were from Sigma. Lazaroid U-83836E (LAZ) was from Abcam. Formaldehyde (FA) 16% aqueous solution was from Electron Microscopy Sciences. Immunofluorescence and western blots primary antibodies used are anti-H3K9me$_3$ (Millipore 07-442), anti-H3K27Acetyl (Millipore MAS23516), anti-H3Acetyl (Upstate 06599), anti-TRF1 (a courtesy of D. Umlauf, CBI, UT, Toulouse), anti-PML bodies (Genetex 55751), anti-SUZ12 (Abcam D39F6), anti-Lamin B1 (Arigo 67137), anti-alpha-tubulin (Sigma T6199), anti-E-Cadherin (BD Biosciences 610181), anti-EpCam (Santa Cruz 25308), anti-beta-Actin (Santa Cruz 69879), anti-GAPDH (Invitrogen 2C2) and AlexaFluor™ Plus 647 Phalloidin (Actin, Invitrogen). Secondary antibodies labeled with Alexa Fluor™ 555 and 647 (Thermo Fisher Scientific) were used for fluorescent imaging while anti-rabbit-HRP and anti-mouse-HRP (GeneTex) were used for western blotting.

**Culture of adherent human cells.**

The human cell line Hep3B was obtained from ATCC® (HB-8064), the human cell lines T24, PC3 and C4-2B were a courtesy of Pr. Heinz Gornitzka team (LCC, CNRS, Toulouse), the human cell line WI-38 stably transfected by ER-GFP-Raf1 and induced 3 days into senescent by adding 4-hydroxytamoxifen, was provided by Pr. Carl Mann [3]. Hep3B, T24, PC3, C4-2b cells were cultured at 37 °C, 5% $CO_2$ and 21% $O_2$, while WI-38 cells were cultured at 37 °C, in normoxia (5% $O_2$). Hep3B cells were cultured in DMEM medium (Gibco), WI-38 cells were cultured in MEM (Gibco) medium while the other cell lines were cultured in DMEM/F-12 (Gibco) medium, all supplemented with 10% FBS (Biosera), L-Glutamine (Gibco), Na-Pyruvate (Gibco), NEAA (Gibco). Cells' subculturing was obtained with trypsin-EDTA 0.25% (Gibco).

**Equipments.**

**Widefield imaging.** For fluorescence imaging a widefield Nikon Ti eclipse microscope equipped with a 20x/0.8 oil and 100x/1.4 oil objectives, Hamamatsu OrcaFlash4v2 camera, an Okolab temperature/$CO_2$ control module and NIS element software was used. For excitation

Lumencor LED system with 390/22nm, 475/34nm, 542/33nm, 575/33nm and 628/40nm filters was used depending on fluorophore. Semrock emission filters 445/20nm, 536/40nm, 539/40, 641/29 and 676/29 were used.

**RIM imaging.** 3D was performed on cells expressing using an upgrade of the system and method described previously [4]. In brief, 3D images were acquired during using an inverted microscope (TEi Nikon) equipped with a ×60 magnification, 1.35 NA objective (CFI Plan Apochromat Lambda S 60XC Sil NIKON) and SCMOS camera (ORCA-Fusion, Hamamatsu). A commercial acquisition software (INSCOPER SA) enables a whole-cell single-timepoint 3DRIM acquisition in only 6 s under a low-photobleaching regime (1 W $cm^{-2}$). Fast diode lasers (Oxxius) with wavelengths centered at 488 nm and 561nm (LBX-488-200-CSB end LBX-561-200-CSB) were used to produce a TEM00 2.2-mm-diameter beam. The polarization beam was rotated with an angle of 5° before hitting a X4 Beam Expander beam (GBE04-A) and produced a 8.8 mm TEM00 beam. A fast spatial light phase binary modulator (QXGA fourth dimensions) was conjugated to the image plane to create 200 RI by each plane (*14*). 3D image reconstruction was then performed as described previously [4] and at GitHub (https://github.com/teamRIM/tutoRIM).

**Optical Tweezers.** Experiments were performed using a home-built setup dedicated to optical trapping measurements. The system is a combination of a fluorescence microscope and an active optical trapping system. This trapping system has two components, one to precisely control the position of the optical trap on the sample and a back focal plane interferometric (BFPi) system to track the position of the trapped object relative to the center of the laser [5,6]. For calibration and the measurements described below, a steering mirror conjugated to the pupil plane of the microscope (Thorlabs FSM75-P01) is used to achieve nanometric laser deflection. A 3 axes piezo electric stage (Piezo concept BIO3) is also used to precisely focus on the trap object. The optical trapping system is implemented in a conventional inverted fluorescence microscope (Leica DMI6000 B). The lens used is a 100x magnification lens with a numerical aperture of 1.4 (Leica HCX PL APO). To optimize the fill factor of the infrared laser at 85% of the objective lens pupil plane, an X4 afocal telescope consisting of two relay lenses is used (Thorlabs ACA254-050-1064 and ACA254-200-1064). The focused diffraction spot at the objective exit has a width of 600 nm at mid-height. This allows an efficient optical trap. The microscope is optimized for simultaneous GFP and RFP imaging combined with the infrared laser trap thanks to 3 band dichroic (Semrock Di03-R405/488/561/635-t1-25x36). Imaging and optical tweezers are achieved by combining a National Intruments acquisition card with the Inscoper synchronisation box. Inscoper software manages all the synchronisation steps. Regarding calibration and measurements, the first step was to capture an internal lipid droplet (with an index close to 0.5 and an average size of 1 µm) and use it as a probe. The laser used for trapping was centred at 1064 nm and the power injected into the sample was 200 mW. At this power and wavelength, the behaviour of the tissue is not affected.

To measure the tension, a 320nm laser square step with a period of 0.12 Hz was used for back focal plane interferometry tracking. The height of this step remains within the linearity of the position detector and allows us to calibrate the detector repose for each measurement in order to convert V into nm according to the size of the lipid droplet trapped. A Python program then averages 5 relaxation x curves for each measurement to reduce thermal or ATP fluctuations.

The fluctuation dissipation theory optical tweezers (FDT optical tweezers) force calibration method has been implemented to simultaneously calibrate the laser trap and the tracker conversion coefficient (V/nm) [7,8]. In this method, when the properties of the optical medium are unknown or heterogeneous, the fluctuation dissipation theory at high frequency is used to calibrate the optical tweezers. A passive and active trajectory consisting of the highest frequency at 1HZ is recorded for 17 seconds to calibrate the optical trap for each measurement. The k-, beta- and elastic response of the medium are estimated simultaneously with an adapted program. The program is inspired by a multiplexed FDT method [9]:

The stiffness of the optical tweezers is estimated with an absolute error of 3% using the following expression from the fluctuation dissipation theorem [9], which takes into account all the harmonics from the square excitation.

$$k_t(n\omega_d)=\sum_{n=1,3,5...}^{\infty} -2k_b\,T Re\left[\frac{P_{active}}{P_{passive}}\right]$$

With $P_{active}$ the power active spectrum end $P_{passive}$ the power passive spectrum. The high-frequency analysis and the number of harmonics provide a very robust estimation of $k_t$ . The mechanical response of the medium can be reconstructed independently at each harmonic frequency once $k_t$ is known. The complex susceptibility of the trapped droplet is obtained from the passive fluctuations as follows for each harmonic frequency $\omega=n\omega_d$

$$\chi(\omega)=\frac{\langle|x(\omega)|^2\rangle}{2k_BT}$$

The inverse susceptibility can then be written as:

$$\chi^{-1}(\omega)=k_t+6\pi RG^*(\omega)$$

where $G^*(\omega)=G'(\omega)+iG''(\omega)$is the effective complex viscoelastic modulus of the surrounding medium probed at frequency $\omega$ and $R$ is the radius of the trapped droplet. Therefore, the complex modulus is directly reconstructed for each odd harmonic of the square-wave drive.

$$G^*(n\omega_d)=\frac{1}{6\pi R}[\chi^{-1}(n\omega_d)-k_t]$$

This multi-harmonic reconstruction method provides independent estimates of G′(ω) and G″(ω) over a broad frequency range. Furthermore, the dispersion of the reconstructed values across harmonics provides a direct estimate of experimental uncertainty and mechanical heterogeneity. A phenomenological viscoelastic model (e.g. power-law or Kelvin–Voigt behaviour) can subsequently be fitted to G′(ω) and G″(ω) when needed. The following equation, which comes from 9, has been used to fit a mixture of cytosol and actin medium, and it has proven to be very effective:

$$G^*(\omega)=\frac{1}{6\pi R}\left[k_{m0}+\frac{k_{m1}(i\omega)^{\alpha}}{\Gamma(\alpha)}+i\omega\gamma\right]$$

**Photoactivation and FRAP microscopy.** Laser fluorescence recovery after photobleaching (FRAP) experiments were performed using a diode laser (488 nm) coupled to a galvanometer-based scanning system (ILAS2, Roper Scientific), which was mounted on an inverted Leica DMI6000B microscope. The laser beam was focused through a high-numerical-aperture oil-immersion objective (Plan-Apochromat ×100/0.7–1.4, Leica) and nuclear bleaching was performed in the focal plane along a 4 µm line at the centre of the nucleus during 80 ms at 100% power. Live imaging was performed using a wide-field microscope equipped with a cooled CCD camera (HQ2, Roper Scientific; 64.5 nm/pixel), a GFP filter and HBO illumination to limit photobleaching. Image acquisition was controlled by Metamorph software coupled to ILAS, with images captured every 0.5 s for 7.5 s prior to ablation and for 31.5 s following bleaching.

**Optofixation methods.**

**Wide-field cell islet irradiation.** 35 mm culture micro-dishes (Ibidi®) containing monolayers of adherent human cell lines (Hep3B, PC3, T24, C4-2B, WI-38h) were incubated in the dark (previous conditions), with nuclear trackers in standard conditions (DAPI, Hoechst 33258, Hoechst 33342, CPG, DRAQ-5, NucRed™ live 647) or at 10 µM in basic MEM without phenol red supplemented with Na-pyruvate and NEAA, at 37 °C for 4-24h (PAL, BER, JAT, CRY, NMC). PAL-treated Hep3B cells were irradiated using wide-field microscope (475/34 nm with a range 3.5 to 15 W/cm$^2$ from outer limit to center of the irradiation beam). For others dyes optofixing assays, the same conditions were applied with 390/22nm excitation Lumencor LED for DAPI, Hoechst 33258 and Hoechst 33342; 475/34nm excitation for CytoPainter Green, Berberine chloride, jatrorrhizine chloride, cryptolepine iodide, N-methylcryptolepine iodide; and 628/40nm excitation for DRAQ-5, NucRed™ live 647. Images were acquired using a NIS Element software.

**Wide-field SC irradiation**. Laser irradiation (488 nm, 11 kW/cm²) was applied to SC, laser beam focused on one nucleus thanks to ASTER illumination system from Abbelight™ SAFe MN 180. Irradiation was performed during 7.5 s and time-lapse BF imaging of 300 frames (exposure time 200 ms – no delay between time points) before and after irradiation.

**Portable LED irradiation.** A 60 mm culture micro-dish containing Hep3B cells as a monolayer (*ca.* $5.10^6$ cells) was treated with PAL (previous conditions) then irradiated for 15 min on a bed of ice using a 467 nm portable PR160L lamp (Kessil®, USA) set at 100 % intensity, positioned directly on the open dish (final T° = 13 °C).

**Small-molecule co-labeling**. Prior or following optofixation, PAL-treated (previous conditions) Hep3B cells were labelled with MitoRed, DAPI, Hoechst 33342, propidium iodide, 7-AAD, DRAQ-5, SYTOX™ Blue, NucRed™ live 647 or NucView® 530 Red Caspase-3 Dye following the supplier instructions. Wide-field fluorescence imaging was performed using the proper acquisition channels for every dye.

**Time-lapse PAL intensity measurements.**

Wide-field images using a 100x1.4oil objective of PAL-treated cells, with or without small molecules inhibitors, were analyzed by measuring the fluorescence intensity at each time point (200ms) during a 5 min irradiation. Analyze were performed using ImageJ v1.54p software and data were plotted using PlotTwist [10].

**Assessment of nuclei size distribution.** PAL-treated cells were light-irradiated as previously described. Acquisitions were performed on widefield (description above) using 20x oil objective. Quantifications were performed with ImageJ v1.54p software. In Fig.2, for illuminated PAL-treated cells, two defined regions of interest (ROI) were used to delimited R1 and R2. The same R1-ROI was used in condition of illuminated cells without PAL. Nuclei areas were measured after a thresholding on PAL fluorescence for "PAL + hv" condition or DAPI signal in conditions "hv" and "PAL". In Fig. 3, nuclei areas in region R1 were measured using the same R1-ROI. Violin plot was obtained using the shiny app PlotsOfData [11].

**Phenotypic recapitulation of PAL-mediated optofixation.** PAL-treated (previous conditions) Hep3B cells were exposed to the following aldehydes: FA (4 % in PBS, RT, 10 min), 4-HNE (100 µM – 2 mM in MEM from a 1 M stock solution in ethanol, 37 °C, 15 h), MDA or ACR (100 µM – 1 mM in MEM from pure products, 37 °C, 15 h), GA or MGA (100 µM – 1 mM in MEM from *ca.* 40% stock solutions in water, 37 °C, 15 h). Wide-field imaging was performed at various timepoints to acquire PAL fluorescence (previous conditions) and assess cell fixing.

**Precipitate and SDS-PAGE analysis.** PAL-treated (previous conditions) Hep3B cells were irradiated using a LED lamp (previous conditions). Co-experiments consisted in cells treated or not with PAL (above conditions) then with FA (4% in PBS) incubated at room temperature for 30 min, 4HNE (1 mM in MEM from 100 mM stock in EtOH) or ACR (1 mM in MEM) incubated for 24 h at 37 °C. Control experiments consisted in untreated cells, and cells treated separately with PAL or light (above conditions). The supernatants were discarded, and the cell layers were washed with ice-cold PBS (3 x 200 µL). The cell layers kept on ice were treated with 100 µL of ice-cold dithiothreitol-free lysis buffer containing Tris pH 7.5, 1 % NP-40, β-glycerophosphate (10 mM), sodium fluoride (10 mM), sodium *o*-vanadate (10 mM) and cOmplete protease inhibitor cocktail (Roche), incubated for 1 min, scrapped and transferred into 600 µL Eppendorff tubes for sonication (3 cycles of 10s with 30% of amplitude). Crude cell lysates were frozen at -20 °C overnight then thawed, centrifuged (15,000 rpm, 5 min, 4 °C), kept on ice and the precipitates (i. e. non-soluble parts of crude cell lysates) were separated from the clear supernatants. The precipitates were resuspended in minimal quantities of ice-cold PBS, quantitatively transferred to transparent 250 µL PCR-type Eppendorff tubes and rewashed thrice by resuspension in ice-cold PBS. Observation was performed under UVA light in a CN-15 UV darkroom (Labortechnik). Lysates were dosed for protein content (micro-BCA assay, Thermo Fisher Pierce) (table S2) then deposited (10 µL) on a SurePAGE™ Bis-Tris 4-12 % polyacrylamide precast gel (GenScript). Migration was performed in MES-SDS 1M buffer at 20 °C and 150 V. Gels were visualized on a ChemiDoc scanner (BioRad) using Alexa488, DL488, Cy2, Cy3 and Cy5 fluorescence channels at 5-120 s exposure (showing an absence of signal), then stained by Coomassie Blue and revisualized on the ChemiDoc scanner for image acquisition. The image was processed in black and white for consistency of depiction with the precipitates (i. e. proteins in white).

**Scavenging of lipid aldehydes.** POA (1 mM in MEM, made from a 0.5 M sterile water stock solution) or COS (20 mM in MEM, made from a 20 M sterile water stock solution), were incubated 15h on PAL-treated Hep3B cells (previous conditions) before their irradiation with wide field conditions.

**Assessment of lipid peroxidation.** PAL-treated Hep3B cells (previous conditions) were illuminated in wide-field conditions (as above). BODIPY™ 581/591 $C_{11}$ (1 µM in MEM, made from a 1 mM DMSO stock solution diluted 1000x in pre-warmed MEM by vortexing at maximal speed without interruption for 30 s, 37 °C) was added 30min, 1h or 2h after irradiation. Control experiments consisted in cells just treated with PAL ("PAL") or light ("hv") (above conditions) and fluorescence measurements at identical timepoints. Acquisitions were performed on widefield (description above) using 100x1.4oil objective. Squares region of interest (25 µm²) were placed aleatory in cytoplasm to avoid PAL nucleus signal and measure the integrated density intensities of green (Ex 475/30 nm, Em 536/40 nm) and red (Ex 575/33 nm, Em 641/75 nm) using ImageJ software (details above) for each time points and conditions. Ratiometric (Green/Red) values were plotted as violin plot using the shiny app PlotsOfData [11].

**Inhibition of lipid peroxidation.** PAL-treated (previous conditions) Hep3B cells were exposed to the following antioxidants : sodium ascorbate (1 mM, 15h), *N*-acetyl-L-cysteine (5 mM, 15h), glutathione (1 mM in MEM, 37 °C, 1.5h), gallic acid (10 µM in MEM, 37 °C, 15h), α-tocopherol (100 µM -1 mM in MEM, made from a 1 mM DMSO stock solution diluted in pre-warmed MEM by vortexing at maximal speed without interruption for 30 s, 37 °C, 50 min), LAZ (100 µM in MEM, 37 °C, 50 min). Cells were then light-irradiated using wide-field microscopic irradiation (previous conditions), PAL fluorogenesis and nuclei size were quantified in the illuminated regions (as above).

**Assessment of ROS production.** PAL-treated Hep3B cells (previous conditions) were illuminated by wide field microscope as previously described. Control experiments consisted in non-irradiated PAL-treated cells or irradiated-cells without PAL. Then, ROS Brite™ 670 probe (1 µM, AAT Bioquest) was added 1 h after irradiation during 15 min at 37°C. Cells were then washed with PBS before image acquisitions by wide field microscopy (Ex 628/40 nm, Em 676/29 nm) and measurements were performed as in the "Lipid peroxidation" section. Datas were plotted as violin plot using the shiny app PlotsOfData [11].

**Inhibition of ROS production.** PAL-treated (previous conditions) Hep3B cells were exposed to the ROS inhibitor CEA (6.25 mM in MEM, 37 °C, 1 h) or incubated for 15h in normoxia (5% $O_2$) before being light-irradiated for optofixation by wide-field microscopy (above conditions). PAL fluorogenesis and nuclei size were quantified in the illuminated regions as above.

**Native immunofluorescence labeling**. Following optofixing (above conditions), PAL-treated Hep3B light-irradiated cells were directly incubated with BSA 3% in PBS during 30 min, followed by primary antibodies diluted in BSA 3%-PBS (2h, RT) or phalloidin (1h, RT). Then, cells were washed 3 times and incubated with Alexa Fluor™ antibodies (1:1000) during 1h at RT and washed again 3 times before imaging in PBS. Comparative reference immunolabelling's were performed in Hep3B cells extemporaneously fixed with formaldehyde

(FA, 4% in PBS, RT, 10 min), washed with PBS and then permeabilized 5 min with Triton X-100 0.5% and blocked with 3% BSA in PBS during 30 min. FA fixed and labelled cells were imaged in the mounting solution Citifluor™.

**Statistical analysis.** For Figs. 2E, 3B, 3D, 3E all statistical analysis were performed with a randomized test using the shiny app PlotsOfDifferences [12]. Statistical details of the experiments (data normalization, n or p-values) are indicated in the figure legends.

**Western blot analysis.** Following lysis of PAL-treated optofixed cells for 0, 5, 10 or 15 min using a LED lamp (previous conditions), 40 µg of total proteins (assessed by Micro BCA™ Protein Assay, Thermo Fischer) was prepared with Laemmli buffer following DTT addition and loaded on SDS-Page gels (4-20% acrylamide gradient, BioRad). Proteins were migrated in Tris-glycine buffer and transferred on nitrocellulose membrane with a semi-dry Trans Blot Turbo System (Biorad). Membranes were blocked with PBS containing 0.1% Tween-20 and 5% non-fat dry milk, prior to incubation with primary antibodies, overnight at 4°C. Membranes were then rinsed 3 times with PBS-Tween and incubated with HRP-conjugated secondary antibodies for 1h at RT. Signal was visualized with the enhanced chemiluminescence detection agent (Roche) and the ChemiDoc imaging system (Biorad).

**Hep3B cells transient tranfection**. Hep3B cells (250,000) were seeded in 35 mm grid dishes (Ibidi®), 24h before their transfection with each of the following plasmids (pBos H2B-GFP, pTrip H2B-mCherry, pcDNA3 HA-GFP, pcDNA3 HA-mCherry), using Fugene® HD transfection reagent (Promega). 24h after, transfected cells are light-irradiated using wide-field microscopic irradiation *in situ* (above conditions) at 475/34 nm for GFP overexpressed proteins and at 628/40 nm for mCherry overexpressed proteins.

#### Supplementary Text

##### *Optical Power Deposition and Thermal Diffusion from a Top-Hat Microscope Focus*

The use of focused laser or lamp irradiation through a microscope objective is widely used in fluorescence microscopy, FRAP, photobleaching, optogenetics, and photothermal experiments. We consider the following configuration, with optical power at the entrance pupil of the microscope objective (P). The beam is focused onto the sample as a circular, homogeneous (top-hat) spot, or Gaussian for the Abbe light system.

Unless otherwise stated, we make the following assumptions:
• unit transmission of the objective and optical path;
• negligible reflection losses at the sample surface;
• homogeneous absorption within the illuminated region. These assumptions can be relaxed later by introducing transmission and absorption coefficients.

The illuminated area seen through the microscope objective in the object plane is:

$$A = \pi r^2$$

For a top-hat intensity distribution, the surface power density (irradiance) is uniform and given by

$$I = \frac{P}{A}$$

This corresponding to 11,3 kW.cm$^{-2}$. The total optical energy incident during the exposure is

$$E = P, t_{exp}$$

If only a fraction A of the incident light is absorbed, the absorbed energy becomes

$$E_{abs}=A_{abs}, E.$$

The absorption coefficient depends on the wavelength and the optical properties of the sample. The Power absorbed in the sample is :

$$P_{abs}=A_{abs} I \pi r^2$$

The temperature field T(r,t) is governed by the heat equation.

$$\rho c \frac{\partial T}{\partial t}=k \nabla^2 T+Q(r,t),$$

where:

ρ is the mass density (kg $m^{-3}$)
c is the specific heat capacity (J $kg^{-1}.K^{-1}$),
k is the thermal conductivity (W $m^{-1}$ $K^{-1}$), for water the value is 0.6 W $m^{-1}K^{-1}$
Q is the volumetric heat source (W $m^{-3}$).

Introducing the thermal diffusivity

$$\alpha=\frac{k}{\rho c}$$

We obtain,

$$\frac{\partial T}{\partial t}=\alpha \nabla^2 T+\frac{Q}{\rho c}$$

The absorbed surface power density is

$$I_{abs}=A_{abs} I$$

If absorption occurs over an effective depth h, the corresponding volumetric heat source is

$$Q=\frac{I_{abs}}{h}$$

For a top hat beam

$$Q=\frac{A_{abs} P}{\pi r^2 h}, for\ |r| \leq r,$$

The characteristic diffusion length after time t is

$$l_{th}(t)=\sqrt{4\alpha t}.$$

For water-like media α =10^-7 $m^2$ s-and dt = 7 s, this yields diffusion lengths of several tens of micrometers, The approximate temperature increase is

$$\Delta T \sim \frac{P_{abs}}{4\pi k l_{th}}$$

The following table resumes the power density and the estimation of the increase of temperature for each experimental conditions :

| **Case** | **Spot diameter** | **Exposure time** | **Irradiance** | **Thermal diffusion length ($l_{th}$ )** | **Temperature rise (ΔT )** |
|---|---|---|---|---|---|
| Diffraction-limited | ≈ 1 µm | 200 ms | 7.5 kW/cm² | ≈ 11 µm | 0.02–0.05 °C |

| Case | Spot diameter | Exposure time | Irradiance | Thermal diffusion length ($l_{th}$ ) | Temperature rise ($\Delta T$ ) |
|---|---|---|---|---|---|
| Small spot | 15 µm | 7 s | 11 kW/cm² | ≈ 2 mm | 0.1–0.3 °C |
| Medium spot | 50 µm | 1 min | 250 W/cm² | ≈ 6.5 mm | 0.1–0.3 °C |
| Very large spot | 300 µm | 5 min | 3.5-10 W/cm² | ≈ 13 mm | 0.1–1 °C |

***Unified extraction of transport and optofixation kinetics***

The table below summarizes the quantitative parameters extracted from a combination of photoactivation, fluorescence recovery after photobleaching (FRAP), optical tweezers and label-free imaging experiments, all of which were performed under controlled irradiation conditions. While these measurements probe different physical properties and cellular compartments, they converge towards a unified description of a light-induced optofixation process governed by a few kinetic rates. Taken together, these results demonstrate that optofixation is a continuous, power-dependent process that progressively suppresses molecular diffusion, enhances viscoelastic rigidity and ultimately results in a solid-like cellular state. The consistency of the extracted kinetic parameters across fluorescence-based, mechanical and label-free measurements strongly suggests that a unified ROS-driven mechanism underlies the optofixation transition.

| Phase / Method | Observable | Physical quantity probed | Model / Assumption | Typical timescale | Estimated parameter |
|---|---|---|---|---|---|
| **BF – lipid droplet tracking** | Arrest of droplet motion | Cytosolic transport | $(D_{\text{eff}}(t)=D_0 e^{-k_{\text{fix}}t})$ | ~2 s at 11 kW/cm² | ($k_{\text{fix}}$ ~ 0.35 $s^{-1}$) |
| **Optical tweezers (cytosol)** | Elastic modulus increase (×12) | Cytosolic stiffening | $(G'(t)=G'0e^{k\text{fix}t})$ | ~15 s at 40 W/cm² | ($k_{\text{fix}}$ ~ 0.15–0.2 $s^{-1}$) |
| **Phase I – Photoactivation (low power)** | Fluorescence rise at fixed distance | Molecular diffusion + photoactivation | Diffusion with photoactivation rate ($k_{\text{pa}}$) | Plateau reached in ~30 s | ($k_{\text{pa}}$ ~ 0.3~$s^{-1}$) |
| | Early signal appearance (~3 s) | Initial spatial spreading | Gaussian broadening ($\sigma^2(t)=2Dt$) | Detectable in ~3 s | ($\sigma$ ~ 4.5 $\mu$m), ($D$ ~ 4.6 $\mu m^2.s^{-1}$) |
| | Spatial profile of activated fluorescence | Diffusion coefficient | Free diffusion approximation | 3–30 s | ($\sigma$ ~ 0.8–1.0~$\mu$m), ($D$ ~ 0.014 $\mu m^2.s^{-1}$) |
| **Phase II – Mixed PA + optofixation** | Decay of effective diffusion | Progressive immobilization | $(D(t)=D_0 e^{-k_{\text{fix}}t})$ | 10–50 s at 5 W/cm² | ($k_{\text{fix}}$ ~ 0.03–0.1~$s^{-1}$) |
| | Growth saturation of ($\sigma$^2($t$)) | Arrest of spatial spreading | Diffusion + fixation + bleaching | 10–50 s | Consistent with ($k_{\text{fix}}$) above |
| | Fluorescence intensity dynamics | PA + bleaching + fixation | Reaction–diffusion model | — | ($k_{\text{bleach}}$) small at low power |
| **Phase III – FRAP (high fixation)** | Fluorescence recovery plateau | Mobile fraction | Reaction–diffusion with fixation | 300 s | Mobile fraction ($f_m \ll$ 1) |
| | FRAP recovery time | Effective diffusion of mobile pool | Corrected for photoactivation | — | ($D_{\text{eff}}$) strongly reduced aroud 0.0001 $\mu m^2.\ s^{-1}$ |

| Phase / Method | Observable | Physical quantity probed | Model / Assumption | Typical timescale | Estimated parameter |
|---|---|---|---|---|---|
| **BF – lipid droplet tracking** | Arrest of droplet motion | Cytosolic transport | $(D_{\mathrm{eff}}(t)=D_0 e^{-k_{\mathrm{fix}} t})$ | ~2 s at 11 kW/cm² | $(k_{\mathrm{fix}} \sim 0.35\ \mathrm{s}^{-1})$ |
| **Optical tweezers (cytosol)** | Elastic modulus increase (×12) | Cytosolic stiffening | $(G'(t)=G'0 e^{k\mathrm{fix}t})$ | ~15 s at 40 W/cm² | $(k_{\mathrm{fix}} \sim 0.15\text{–}0.2\ \mathrm{s}^{-1})$ |
| | Comparison with $(\sigma^2)$ from phases I–II | Consistency of transport arrest | Same fixation rate | — | Unified $(k_{\mathrm{fix}})$ |

***Estimation of the Fixation Rate from BF Dynamics Arrest***

In order to characterize the kinetics of optofixation independently of fluorescence-based measurements, we analyzed the arrest of intracellular and cellular-scale dynamics using time-resolved BF imaging under high irradiation intensity.

Experiments were performed at an irradiation power density of

$$I \simeq 11\text{~kW/cm}^2$$

The motion of lipid droplets was tracked over time, both prior to and during irradiation. In the absence of fixation, their motion can be accurately modelled as an effective diffusion process involving mean squared displacement.

$$\langle \Delta r^2(t) \rangle = 2 d D_{\mathrm{eff}} t$$

where d is the dimensionality and $D_{\mathrm{eff}}$ the effective cytosolic diffusion coefficient. During irradiation, a rapid decrease of droplet mobility was observed, corresponding to a progressive reduction of the accessible mean free path. We model this arrest as an exponential decay of the effective diffusion coefficient:

$$D_{\mathrm{eff}}(t)=D_0 e^{-k_{\mathrm{fix}} t}$$

Where $k_{\mathrm{fix}}$ is the optofixation rate.

***Estimation of the Cytosolic Fixation Rate from Optical Tweezers Microrheology***

To independently quantify the kinetics of cytosolic optofixation, we performed active microrheology measurements using optical tweezers during irradiation (see Optical Tweezers section).

A lipid vesicle of radius $[R \simeq 400\text{~nm}]$ was embedded in the cytoplasm and mechanically trapped. The local complex viscoelastic modulus $G^*(\omega,t)=G'(\omega,t)+iG''(\omega,t)$ , was extracted during continuous irradiation inducing photofixation.

we observed a rapid and pronounced increase of the elastic component of the viscoelastic modulus. Specifically, within $t \simeq 15\text{~s}$ the elastic modulus increased by approximately a factor of 12:

$$\frac{G'(t=15\sim\mathrm{s})}{G'(t=0)} \approx 12.$$

This strong stiffening indicates a rapid transition of the cytosol toward a more elastic, solid-like state during irradiation. We assume that cytosolic stiffening originates from the formation of light-induced crosslinks between cytoskeletal and cytosolic proteins, mediated by reactive oxygen species (ROS). In the regime of weak to moderate crosslinking, the elastic modulus is expected to scale linearly with the density of crosslinks nxl.

$$G'(t) \propto n_{\mathrm{xl}}(t)$$

We model the formation of crosslinks as a first-order kinetic process:

$$\frac{dn_{\mathrm{xl}}}{dt}=k_{\mathrm{fix}}^{\mathrm{cyto}}(n_{\mathrm{xl}}^{max}-n_{\mathrm{xl}}),$$

where $k_{\mathrm{fix}}^{\mathrm{cyto}}$ is the cytosolic fixation rate and $n_{\mathrm{xl}}^{max}$ the saturation crosslink density.

Solving this equation yields:

$$n_{\mathrm{xl}}(t)=n_{\mathrm{xl}}^{max}\left(1\text{-}e^{-k_{\mathrm{fix}}^{\mathrm{cyto}}t}\right)$$

and consequently

$$G'(t)=G'_{max}\left(1\text{-}e^{-k_{\mathrm{fix}}^{\mathrm{cyto}}t}\right)$$

we obtain

$$1\text{-}e^{-k_{\mathrm{fix}}^{\mathrm{cyto}}\cdot 15} \approx 0.95,$$

which leads to

$$k_{\mathrm{fix}}^{\mathrm{cyto}}=-\frac{1}{15}ln(0.05) \approx 0.20\sim\mathrm{s}^{-1}.$$

Taking into account experimental uncertainty, we estimate:

$$k_{\mathrm{fix}}^{\mathrm{cyto}} \sim 0.1\text{-}0.3\sim\mathrm{s}^{-1}$$

***Quantitative photoactivation and FRAP analysis during a viscoelastic optofixation transition in the nucleus***

**1. Summary**

A single laser line and a single wavelength were used for local photoactivation and photobleaching in all experiments. The observation lamp used for fluorescence is disruptive, as it is known to also photoactivate DNA-bound molecules, generate ROS (via these same molecules), gradually induce optofixation, and cause photobleaching. Intensities of the fluorescent signals were measured in areas of interest that had been defined by the user. These areas included a region that had been activated or bleached, as well as the surrounding nuclear regions and a background area. As outlined below, the background noise was eliminated, and the time-dependent fluorescence signals were standardized. To avoid any bias caused by irreversible immobilization, diffusion coefficients were extracted using regime-specific observables. The following table summarizes the regime, observable, method and extraction parameters presented in Figure 2 for each analysis phase.

| Phase | Physical regime | Primary observable | Measurements | Identifiable parameters | What should NOT be done |
|---|---|---|---|---|---|
| **I (t=0)** | Diffusion with weak photoactivation | FRAP recovery | Pure diffusion | D from $\tau_D$ | Negligect photoactivation outside ROI |
| | | $\sigma^2(t)$ | Spatial transport | D (robust) | — |
| | | $F_{out}$ (t) | Photoactivation kinetics | kpa | Interpret as diffusion |
| **II (t=1min)** | Diffusion + photoactivation + photobleaching + optofixation | $\sigma^2(t)$ | Effective transport | Deff ≈D | Extract (D) from intensity |
| | | FRAP recovery amplitude | Mobile fraction | M(t) | Fit $\tau_D$ freely |
| | | $F_{out}(t)$ | Photochemical kinetics | kpa, kbl | Mix transport and chemistry |
| | | FRAP plateau | Immobilization | kfix (indirect) | Assume constant immobile fraction |
| **III (t<5min)** | Optofixation-dominated+ Photoativation+ phpotobleaching | FRAP plateau | Residual mobile fraction | (M) | Extract diffusion coefficient |
| | | Absence of recovery | Complete immobilization | — | Analyze as valid FRAP |
| | | $\sigma^2(t)$ | — | — | Use for diffusion |

#### 2. Phase I: Diffusion after local photoactivation at T=0

*Reaction-diffusion equation*

Following the activation pulse, the evolution of the fluorescent population is governed by

$$\frac{\partial C_f}{\partial t}=D\nabla^2 C_f+k_{\mathrm{pa}}I(r)\left(C_{\mathrm{tot}}-C_f\right),$$

where D is the diffusion coefficient, kpa the photoactivation rate constant, I(r) the local irradiation intensity, and C(tot) the total (activatable) molecular concentration.

*Estimation of the photoactivation rate outside the bleached region.*

Photoactivation occurring outside the bleached region was quantified using a secondary ROI located in a non-bleached, weakly illuminated nuclear area. In this region, diffusion is negligible at short times and fluorescence increase is dominated by local photoactivation.

$$\frac{dF_{\mathrm{out}}}{dt}=k_{\mathrm{pa}}I_{\mathrm{out}}(F_{max}-F_{\mathrm{out}}).$$

The solution of this equation yields

$$F_{\mathrm{out}}(t)=F_{max}\left(1-e^{-k_{\mathrm{pa}}I_{\mathrm{out}}t}\right)$$

from which the photoactivation rate constant kpa was extracted by fitting the initial fluorescence rise. This estimation was performed exclusively in the low-dose regime, ensuring that
photoactivation kinetics were decoupled from diffusion and optofixation.

*Spatial spreading analysis using the second moment*

In addition to intensity-based analysis, diffusion and photoactivation were quantified using the spatial spreading of fluorescence following local photoactivation. The spatial distribution of fluorescence was characterized by its second spatial moment,

$$\sigma^2(t)=\frac{\int |r-r_0|^2\, C_f(r,t)dr}{\int C_f(r,t)\, dr},$$

where r0 denotes the center of the photoactivated region. For a purely diffusive process, the temporal evolution of the second moment follows.

$$\sigma^2(t)=\sigma_0^2+2dDt$$

where $D$ *is* the diffusion coefficient and $\sigma_0$ the initial variance determined by the activation geometry and d is the dimensionality of diffusion (d=2 in the nuclear plane).

This formulation remains valid in the presence of homogeneous low-level photoactivation, as photoactivation affects the total fluorescence amplitude but does not alter the spatial transport term. Consequently, the slope of $\sigma^2$ provides an independent and robust estimate of D, insensitive to photobleaching or intensity normalization. At early times, before significant redistribution of fluorescence from distant regions, diffusion dominates over photoactivation-induced growth, ensuring that $\sigma^2$. Diffusion coefficients were extracted by linear regression of $\sigma^2$ as a function of time, using fluorescence profiles measured perpendicular to the activation boundary.

**3. Phase II : Photoactivation under progressive optofixation**

In phase 2, the dynamics of fluorescence are driven by a combination of diffusion, photoactivation, photobleaching, and ROS-mediated optofixation, all occurring at the same time. This regime is characterized by a transition from classical FRAP dynamics to dynamics dominated by immobilization. At intermediate irradiation doses, the experiment transitions to a mixed regime where photobleaching, diffusion, ROS-induced optofixation, and photoactivation all occur at the same time. The fluorescent molecules are then divided into three distinct populations, which are then separated from each other:

- Cm(r,t):mobile fluorescent molecules,
- Cfix(r,t): immobilized (fixed) fluorescent molecules,
- Cf(r,t) = Cm+ Cfix: total fluorescent population

*Reaction diffusion model:*

The mobile fluorescent population evolves according to

$$\frac{\partial C_m}{\partial t}=D\nabla^2 C_m+k_{\mathrm{pa}}I(r)(C_0-C_m)-k_{\mathrm{fix}}I(r)C_m-k_{\mathrm{bl}}I(r)C_m,$$

where D is the diffusion coefficient, $k_{pa}$ the photoactivation rate, $k_{fix}$ the optofixation rate constant, and $k_{bl}$ the photobleaching rate constant, $I_{(r)}$ is the local irradiation intensity, end C0 the total activable molecule concentration.

The immobilized population grows irreversibly as

$$\frac{\partial C_{\text{fix}}}{\partial t}=k_{\text{fix}}\, I(r) C_m.$$

and is assumed to be non-diffusive on the experimental timescale. Photobleached molecules are assumed to be optically silent and removed from $C_f$.

Fluorescence signal in the bleached region

The measured fluorescence signal within the FRAP region of interest (ROI) is given by

$$F(t) \propto \int_{\text{ROI}} [C_m(r,t)+C_{\text{fix}}(r,t)]\, dr.$$

Photobleaching reduces the total fluorescent population, photoactivation increases it, and optofixation redistributes fluorescence between mobile and immobile fractions.

*Effective FRAP recovery law*

Under the assumption that optofixation occurs on a timescale comparable to or slower than diffusion across the bleached region, the normalized FRAP recovery can be approximated by

$$F(t)=M_t\left(1\text{-}e^{\text{-}t/\tau_D}\right),$$

With

$$\tau_D=\frac{w^2}{4D}$$

where w is the effective width of the rectangular bleached region and M(t) the time dependant fraction mobile. The mobile fraction decreases due to optofixation, photoactivation and photobleaching according to

$$\frac{dM}{dt}=\text{-}(k_{\text{fix}}+k_{\text{bl}})IM+k_{\text{pa}}I(1\text{-}M)$$

At early times, when M(t) varies slowly compared to the diffusion time constant (τ_D), the fluorescence recovery after photobleaching (FRAP) recovery appears quasi-exponential, but with a progressively reduced amplitude. 1-M is the apparent immobile fraction, which is linked to the photoactivation rate k_(pa).

As the diffusion coefficient D is determined independently from the linear growth of $\sigma^2(t)$, the FRAP recovery curve in Phase II does not provide an additional estimate of D. Instead, FRAP recovery constrains the kinetics of the depletion of the mobile fraction.

Deviations from the effective recovery law indicate either the breakdown of homogeneous reaction assumptions or the onset of Phase III, which is dominated by optofixation.

**4. Phase III: Extraction of the Mobile Fraction under Concurrent Photoactivation**

The third phase of the FRAP experiment aims at quantifying the *mobile fraction* of fluorescent molecules within the bleached region. Due to the strong optofixation, the spatial spreading analysis be use. In classical FRAP analysis, this fraction is inferred from the asymptotic fluorescence recovery following bleaching. However, in the present optofixation/photoactivation framework, fluorescence recovery cannot be interpreted solely as diffusion-driven exchange and must be corrected for ongoing photoactivation and photochemical conversion.

*Definition of the Mobile Fraction*

Let I(t) denote the mean fluorescence intensity measured within the bleached region. We decompose the total fluorophore population into a mobile fraction f_m and an immobile fraction f_i = 1 - f_m. In the absence of photoactivation, the asymptotic recovery is classically written as

$$f_m^{\text{classical}} = \frac{I(\infty) - I(0^+)}{I_{\text{pre}} - I(0^+)}$$

where $I_{pre}$ is the pre-bleach fluorescence intensity and $I(0^+)$ the post-bleach intensity. In the present configuration, this definition is no longer sufficient, as fluorescence intensity is continuously modified by light-induced processes during observation.

*Contribution of Photoactivation to Fluorescence Recovery*

Under continuous or repeated irradiation, photoactivation outside the bleached region generates activated fluorophores that subsequently diffuse into the observation area. The fluorescence recovery thus results from two coupled mechanisms:

- diffusive exchange of mobile molecules,
- spatially distributed photoactivation followed by diffusion

The fluorescence signal can therefore be expressed as

$$I(t) = I_{\text{diff}}(t) + I_{\text{pa}}(t)$$

Where $I_{diff}(t)$ corresponds to recovery driven by diffusion of pre-existing fluorescent molecules, and Ipa(t) accounts for newly photoactivated fluorophores entering the bleached region.

Neglecting $I_{pa}(t)$ would lead to an overestimation of the mobile fraction, as part of the recovery is not associated with molecular mobility but with photochemical conversion.

*Corrected Mobile Fraction Estimation*

To isolate the true mobile fraction, the photoactivation contribution must be explicitly removed. Denoting ΔIpa(∞) as the asymptotic fluorescence increase solely due to photoactivation, the corrected mobile fraction is defined as

$$f_m = \frac{I(\infty) - \Delta I_{\text{pa}}(\infty) - I(0^+)}{I_{\text{pre}} - I(0^+)}.$$

The term ΔI_(pa)(∞) can be estimated independently from control experiments without bleaching, or from the spatial spreading analysis introduced in Phase II using the temporal evolution of variance, $\sigma^2$.

*Link with Spatial Spreading Analysis*

As shown in Phase II, photoactivation induces a measurable increase in the spatial variance of fluorescence intensity,

$$\sigma^2(t)=\sigma_0^2+4Dt+\sigma_{\text{pa}}^2(t)$$

where $\sigma_{pa}{}^2$(t) reflects the contribution of newly activated fluorophores. The same term governs the influx of photoactivated molecules into the bleached region, providing a direct, model-consistent correction for the recovery curve.

This coupling ensures that the extraction of the mobile fraction remains physically meaningful and consistent with the underlying reaction--diffusion--photoactivation dynamics.

**5. Clarification about Distinction between mobile fraction and measured recovery signal**

In the context of FRAP experiments performed under continuous irradiation inducing both photoactivation and optofixation, it is essential to distinguish between two related but fundamentally different quantities: the *intrinsic mobile fraction* and the *measured fluorescence recovery signal.*

*Intrinsic mobile fraction.*

The mobile fraction, denoted fm, is a structural and time-independent property of the system once a steady state is reached. It represents the fraction of molecules that remain capable of diffusive motion and are not immobilized by optofixation:

$$f_m=\frac{C_m}{C_m+C_{\text{fix}}},$$

where $C_m$ is the concentration of mobile molecules and $C_{fix}$ is the concentration of optofixed (immobile) molecules. The immobile fraction is defined as fi = 1 - fm.

*Measured recovery signal*

In contrast, the experimentally measured FRAP recovery is a time-dependent observable, denoted M(t), derived from fluorescence intensity measurements within the bleached region:

$$M(t)=\frac{I(t)-I_{\text{bleach}}}{I_{\text{ref}}-I_{\text{bleach}}}$$

where *I(t)* is the fluorescence intensity at time (t), $I_{bleach}$ is the post-bleach intensity, and $I_{ref}$ is the pre-bleach reference intensity.

In standard FRAP experiments without photoactivation or photochemical fixation, the recovery signal can be expressed as

$$M(t)=f_m R(t),$$

where R(t) is the normalized recovery function governed solely by diffusion, with R(0)=0 and R(∞)=1 . In this ideal case,

$$lim_{t\to\infty} M(t)=f_m$$

and the fluorescence plateau directly reports the mobile fraction.

*Effect of photoactivation and optofixation.*

In the present experimental conditions, continuous irradiation simultaneously induces photoactivation of DNA-bound fluorophores and progressive optofixation. As a result, the measured recovery signal includes an additional contribution unrelated to diffusion:

$$M_{\mathrm{meas}}(t)=f_m R(t)+M_{\mathrm{pa}}(t)$$

where Mpa(t) represents the fluorescence increase due to ongoing photoactivation of non-fluorescent or weakly fluorescent molecules.

A minimal expression for this contribution is

$$M_{\mathrm{pa}}(t)=\int_0^t k_{\mathrm{pa}}\, C_{\mathrm{dark}}(t')dt'$$

with $k_{pa}$ the photoactivation rate and $C_{dark}$ the concentration of activatable molecules

*Corrected recovery signal.*

To extract physically meaningful diffusion parameters and the true mobile fraction, the recovery signal must therefore be corrected by the following equation:

$$M_{\mathrm{corr}}(t)=M_{\mathrm{meas}}(t)-M_{\mathrm{pa}}(t).$$

#### Supplementary tables S1-S3, movies S1 & S2, figures S1-S20

**Table S1.** Composition and fluorescent labeling of Hep3B cells by some natural fluorophores

| Compound | Category | Fluorophore type | Labeling (Hep3B cells, MEM, 10-50 µM, 1-15 h) |
|---|---|---|---|
| Guaiazulene | Azulene | Polyene | Negative |
| Piperine | Piperidine alkaloid | Polyene | Negative |
| Atractylodin | Furane polyacetylene | Polyene | Negative |
| Anacardic acid | Salicylic acid polyketide | Salicylic acid | Negative |
| (15:3) Anacardic acid | Salicylic acid polyketide | Salicylic acid | Negative |
| Lasalocid sodium salt | Salicylic acid polyether | Salicylic acid | Negative |
| β-Hydrastine hydrochloride | Isoquinoline alkaloid | *O*-alkylsalicylate | Negative |
| Kynurenic acid | Quinoline-2-carboxylic acid | 4-Hydroxyquinoline | Negative |
| Xanthurenic acid | Quinoline-2-carboxylic acid | 4-Hydroxyquinoline | Negative |

| | | | |
|---|---|---|---|
| Alternariol | Benzocoumarin | 7-Hydroxycoumarin | Negative |
| Alternariol monomethyl ether | Benzocoumarin | 7-Hydroxycoumarin | Negative |
| Coumestrol | Benzofurocoumarin | 7-Hydroxycoumarin | Negative |
| (+/-)-Heraclenin | Furocoumarin | 7-Alkoxycoumarin | Negative |
| Oxypeucedanin | Furocoumarin | 7-Alkoxycoumarin | Negative |
| Byakangelicol | Furocoumarin | 7-Alkoxycoumarin | Negative |
| Byakangelicin | Furocoumarin | 7-Alkoxycoumarin | Negative |
| Colladin | Coumarin sesquiterpene | 7-Alkoxycoumarin | Negative |
| Farnesiferol A | Coumarin sesquiterpene | 7-Alkoxycoumarin | Negative |
| Farnesiferol C | Coumarin sesquiterpene | 7-Alkoxycoumarin | Negative |
| Rutaecarpine | Tryptamine alkaloid | 2-Indoloquinazolinone | Negative |
| Evodiamine | Indoloquinazoline alkaloid | 2-Indoloquinazolinone | Negative |
| Tryptanthrin | Indoloquinazoline alkaloid | Indoquinazolinone | Negative |
| Palmatine chloride | Protoberberine alkaloid | 2-Arylisoquinolinium | Positive (this study) |

**Table S2**. Soluble protein quantification of PAL-treated optofixed cells, in comparison to various negative (i. e., cells exposed to PAL or light or receiving no treatment) or positive (i. e., cells exposed or not to PAL then 4 % FA, 1 mM 4HNE or 1 mM ACR) controls. Results shown are from three independent experiments.

| **PAL** | - | + | - | + | - | + | - | + | - | + |
|---|---|---|---|---|---|---|---|---|---|---|
| **Aldehyde** | - | - | FA | FA | 4HNE | 4HNE | ACR | ACR | - | - |
| **LED irradiation @ 467 nm (15 min)** | - | - | - | - | - | - | - | - | + | + |
| **[protein] (mg/mL)** | 1.815 ± 0.013 | 1.513 ± 0.018 | 1.322 ± 0.013 | 1.082 ± 0.005 | 0.085 ± 0.002 | 0.078 ± 0.001 | 0.262 ± 0.006 | 0.324 ± 0.010 | 1.632 ± 0.018 | 0.940 ± 0.013 |

**Table S3**. Categorization of various cell imaging trackers according to their excitation wavelength, localization in live cells and optofixing capabilities by WF microscopy.

| Dye | Excitation wavelength | Main target organelle | Optofixation |
|---|---|---|---|
| DAPI | 390/22 nm | Nucleus | + |
| Hoechst 33258 | | Nucleus | + |
| Hoechst 33342 | | Nucleus | + |
| SYTOX™ Blue | | Nucleus | - |
| ER-Tracker™ Blue-White | | Endoplasmic reticulum | - |
| Quinine | | Lysosome | - |
| Palmatine (PAL) | 475/34 nm | Nucleus | + |
| Berberine (BER) | | Nucleus | + |
| Jatrorrhizine (JAT) | | Nucleus | + |
| Cryptolepine (CRY) | | Nucleus | + |
| *N*-Methylcryptolepine (NMC) | | Nucleus | + |
| CytoPainter™ Green (CPG) | | Whole cell | + |
| MitoTracker® Green | | Mitochondria | - |
| Papaverine | | Cytosol | - |
| Propidium iodide (PI) | 575/33 nm | Nucleus | - |
| MitoRed® | | Mitochondria | - |
| Nile Red | | Plasmic membrane | - |
| 7-Aminoactinomycin (7-AAD) | 628/40 nm | Nucleus | - |
| DRAQ5™ | | Nucleus | + |
| NucRed™ Live 647 | | Nucleus | + |
| LysoTracker™ Deep Red | | Lysosome | - |
| Verteporfine | | Cytosol | - |

**Movie. S1.** SMLM fluorescence imaging of the time-dependent, irradiation-mediated (488 nm, 11 kW/cm$^2$, 7.5 s) nuclear fluorogenic labeling of PAL-treated SC. Results are representative of three independent experiments.

**Movie. S2.** WF fluorescence imaging of the time-dependent, irradiation-mediated (475/34 nm, 3.5 to 15 W/cm$^2$ from outer limit to center of the irradiation beam, 5 min) nuclear fluorogenic labeling of PAL-treated cells. Results are representative of six independent experiments.

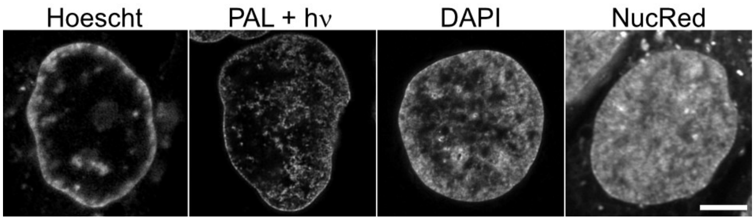


**Fig S1.** RIM images of chromatin labeling after optofixation (PAL+hv) compared to Hoechst 33342, DAPI or NucRed™ live 647 staining (scale bars: 2 µm).

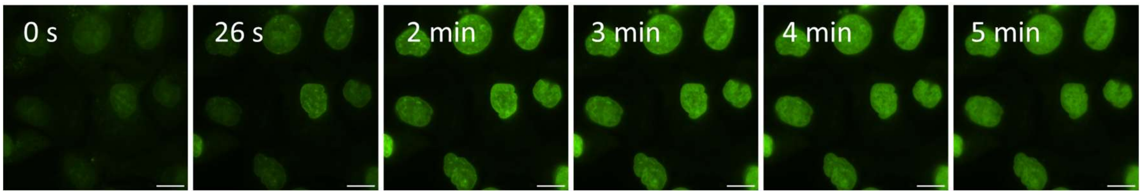


**Fig S2.** Time-lapse WF imaging irradiation of PAL-treated cells after FA fixation, showing progressive nuclear fluorogenesis (scale bar: 10 µm). Results are representative of six independent experiments.

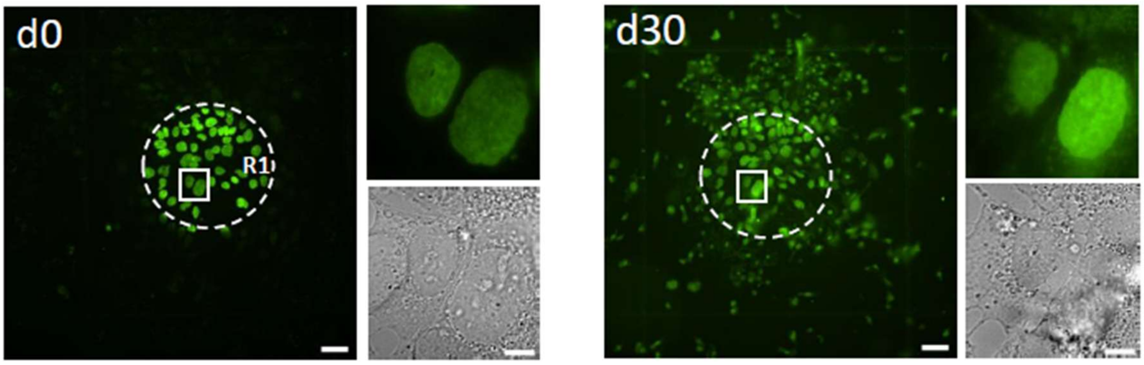


**Fig S3.** Observation of PAL-treated cells immediately after irradiation (*left*, d0) showing persistence more than 30 days (*right*, d30). Crops in R1 region (dotted lines) show nuclear PAL fluorescence and BF images (scale bars: 50 µm; 10 µm in crops). Results are representative of six independent experiments.

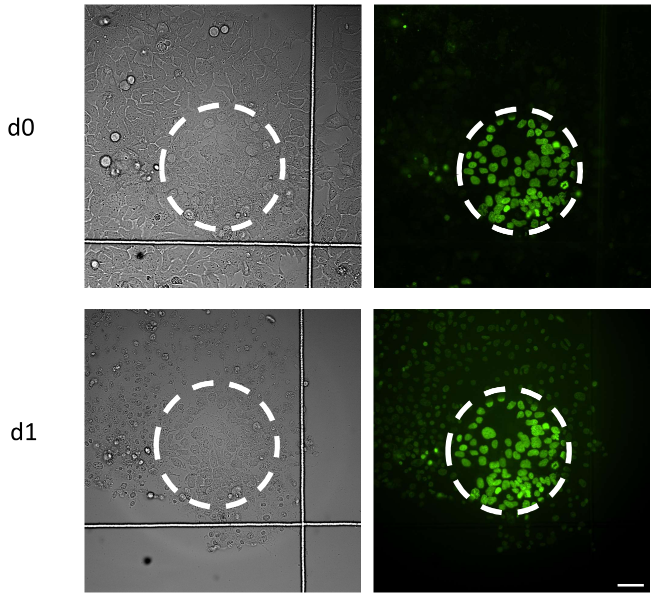


**Fig S4.** Observation of PAL-treated cells immediately after irradiation (*up*, d0) showing resistance after 24 h of triton X-100 treatment (*down*, d1). R1 region is shown by dotted lines (scale bar: 50 µm). Results are representative of three independent experiments.

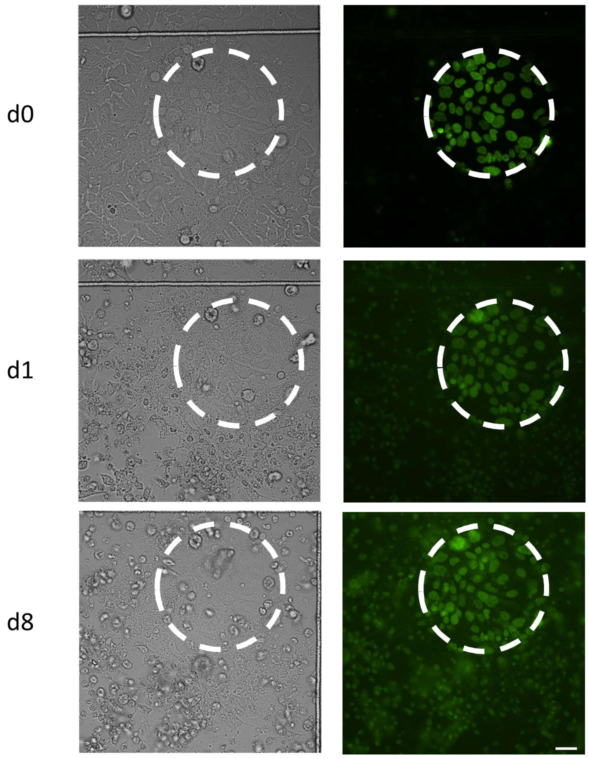


**Fig S5.** Observation of PAL-treated cells immediately after irradiation (*up*, d0) showing resistance to doxorubicine 24 h (*middle*, d1) or 8 days (*down*, d8) after treatment. R1 region is shown by dotted lines (scale bar: 50 µm). Results are representative of three independent experiments.

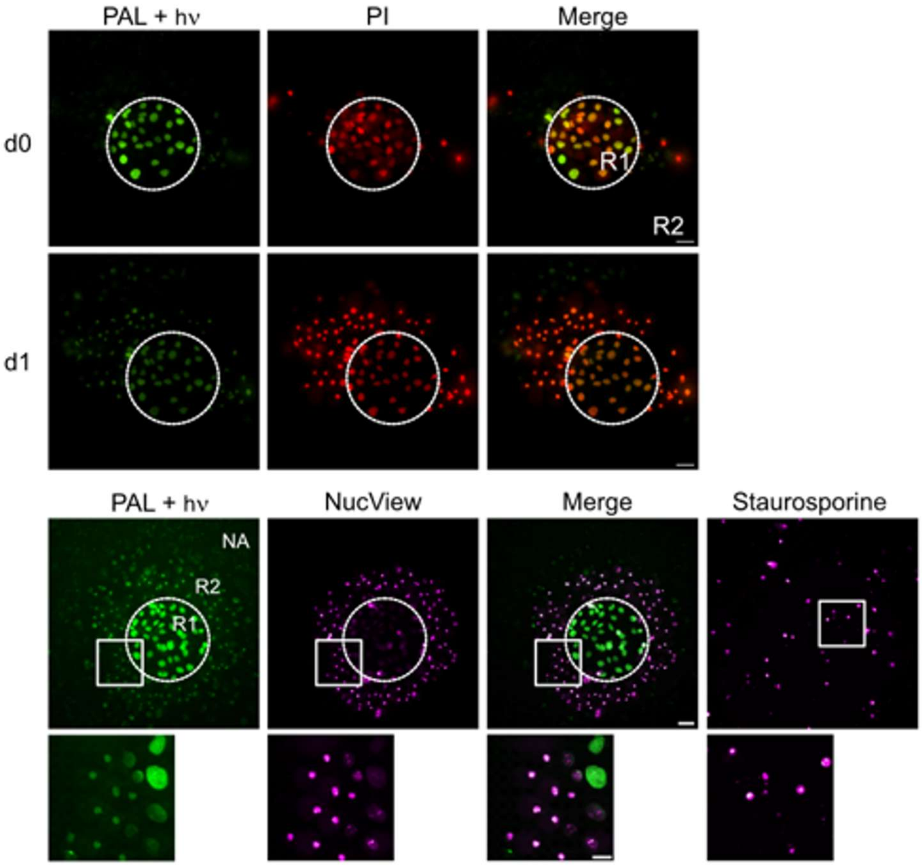


**Fig S6.** Co-staining of PAL-treated cells (green) with propidium iodide (PI, red) or NucView® 530 Red Caspase-3 dye (NucView, magenta). *Top:* PI stains R1 cells (dotted lines), showing a permeabilized status immediately post-irradiation (d0). After 24 h, R2 cells show substantial delayed PI staining demonstrating a compromised state. *Bottom:* NucView faintly stains R1 cells 24 h post-irradiation, showing strong signal in compromised R2 cells. The apoptosis inducer staurosporine is used as a positive control.

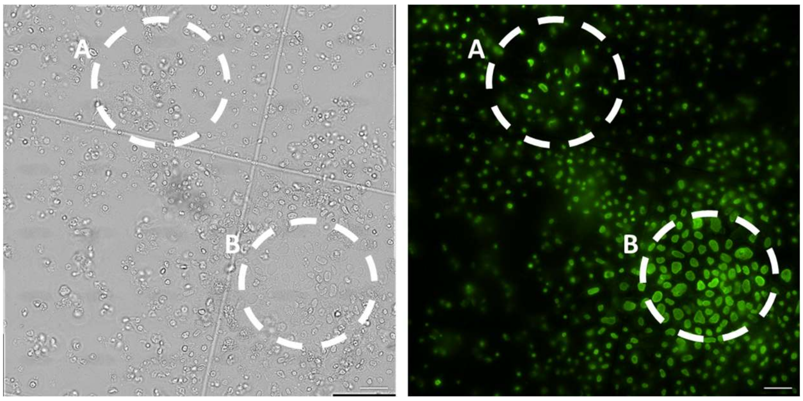


**Fig S7.** Irradiation-dependent fixing state of PAL-treated cells is not recapitulated by sequential induction. In zone A, cells are first irradiated and then incubated with PAL while in zone B, cells are irradiated in presence of PAL (observed after 8 days). Results are representative of three independent experiments.

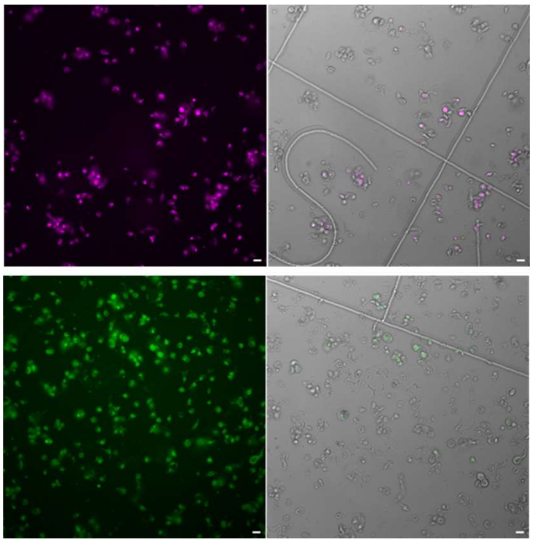

**Fig S8.** Doxorubicine treatment of cells followed by 7AAD labeling (*up*) or PAL labelling (*down*). Results are representative of three independent experiments.

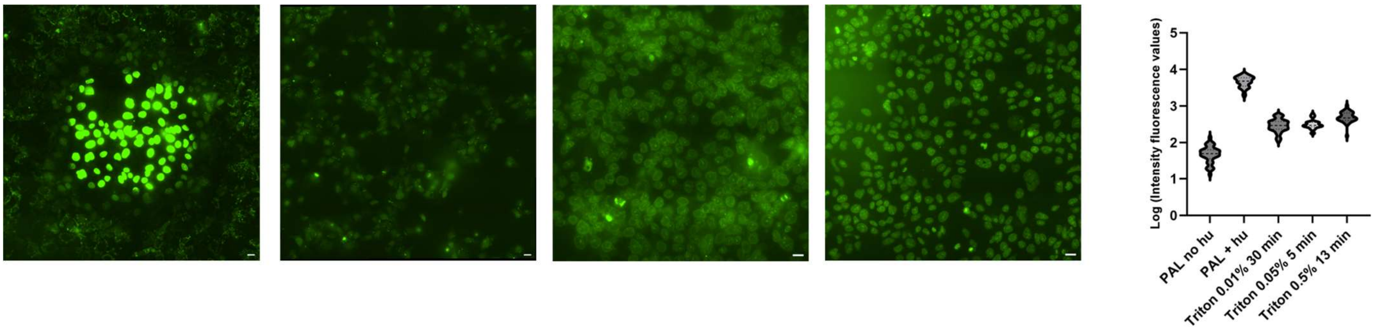


**Fig S9.** PAL fluorescence of cells extemporaneously treated with triton X-100, compared to PAL-light mediated fluorescence (*left*); 0.01% triton 30 min (*middle left*), 0.05% triton 5 min (*middle right*), triton 0.5% 13 min (*right*) and nuclear fluorescence quantification (n=40).

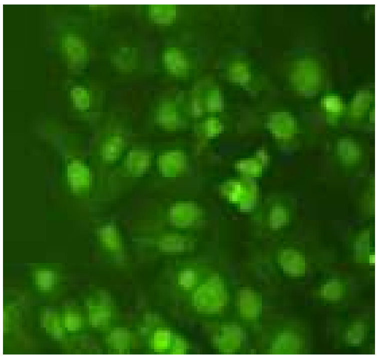

**Fig S10.** PAL fluorescence in FA fixed cells. Results are representative of six independent experiments.

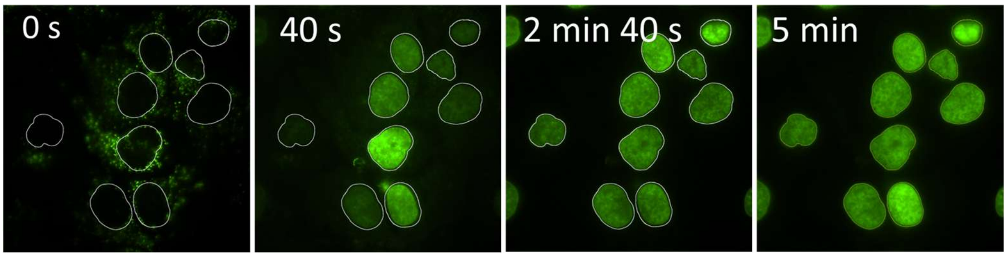


**Fig. S11**-Nuclei area tracking during time-lapse irradiation of PAL-treated cells. Results are representative of six independent experiments.

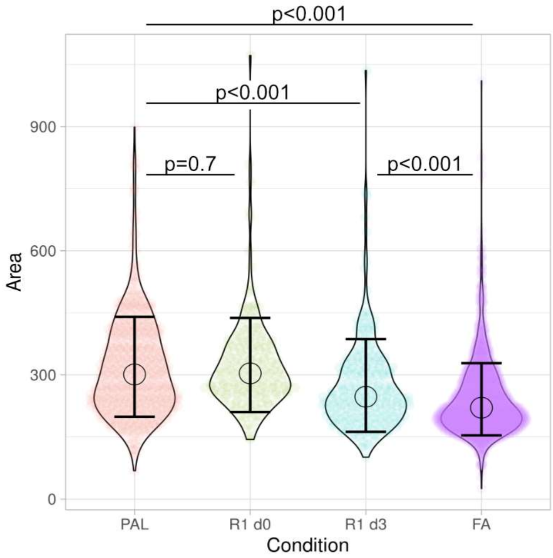


**Fig S12.** Quantification of nuclei areas immediately and 3 days after PAL-mediated optofixation of cells compared to live and FA fixed cell.

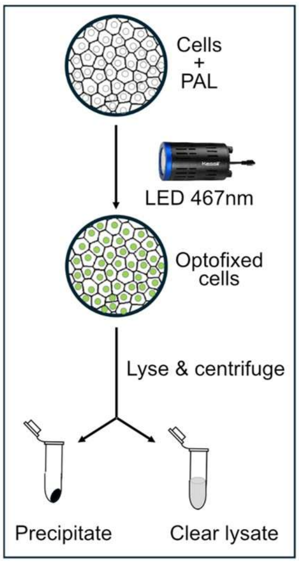


**Fig. S13.** Optofixing procedure of a whole Hep3B cell population using a portable LED lamp towards biochemical analysis.

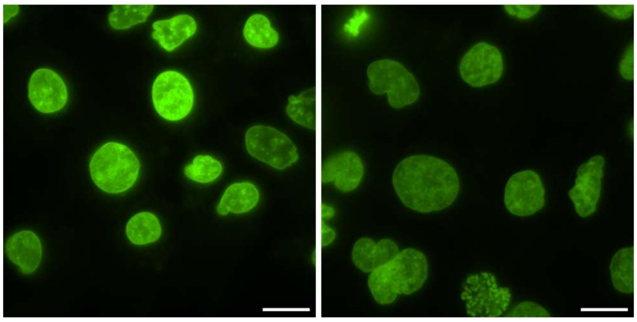

**Fig S14.** PAL-mediated irradiation by a portable LED lamp (*left*), compared to PAL-mediated irradiation by WF irradiation (*right*) (scale bar: 20 µm). Results are representative of six independent experiments.

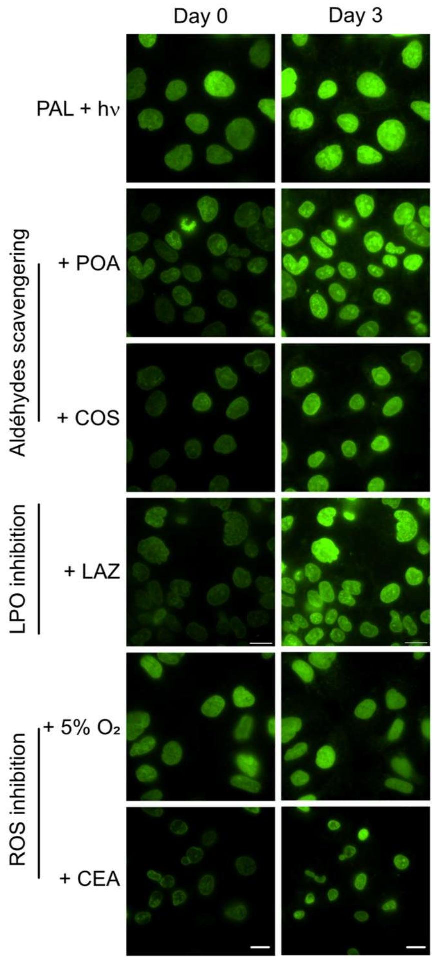


**Fig. S15.** WF imaging of PAL nuclear fluorogenesis at day 0 after irradiation by a WF microscope in the presence of inhibitors (scale bar: 10µm).

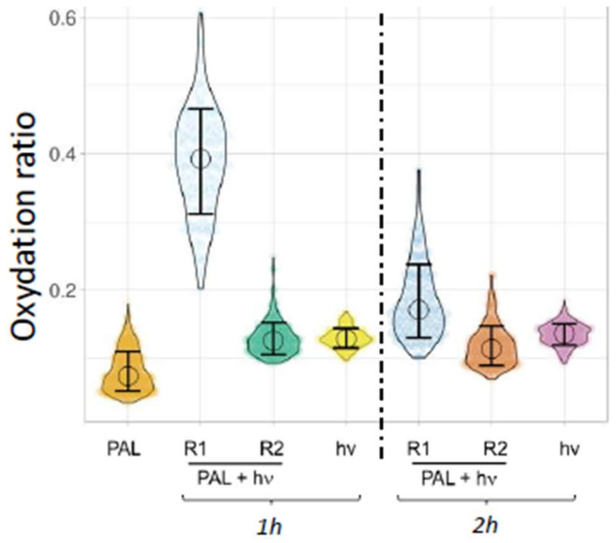


**Fig S16.** Lipid peroxidation (LPO)-quantification in PAL-treated cells using BODIPY™ 581/591-C11 probe 1h or 2h after irradiation of PAL-treated cells (R1, R2) compared to irradiated cells in absence of PAL (hv) and cells incubated with PAL without irradiation (PAL) (n>100 per conditions).

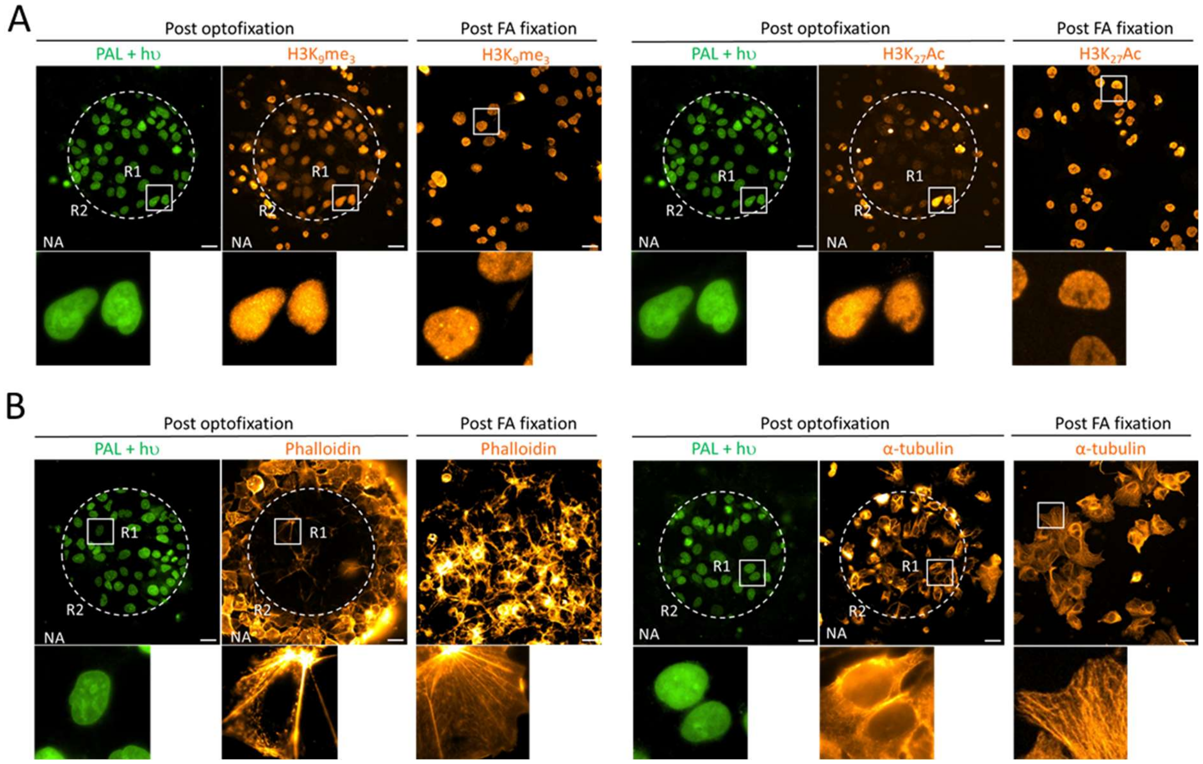


**Fig S17.** Each tested antigen shows: *left* : PAL staining after optofixation (green, dotted circles R1 cells); *middle* : native IF (hot orange, "post-optofixation") compared to *right* : standard IF (permeabilized FA-fixed cells, hot orange,"Post-FA fixation"). Scale bar: 30 µm. **(A)** Nuclear antigens labelling. **(B)** Cytoskeleton antigens labelling. Results are representative of four independent experiments.

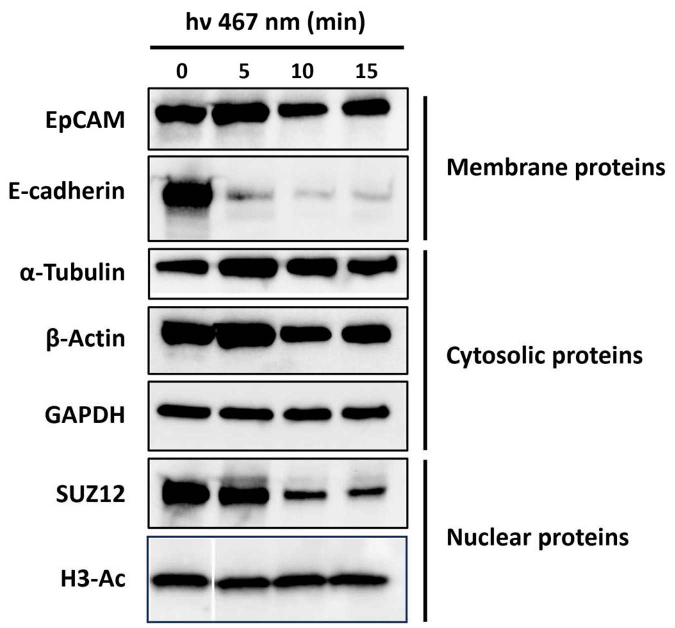


**Fig S18.** Western blot analysis of some antigens after PAL-treated cells optofixation with portable LED lamp. Results are representative of three independent experiments.

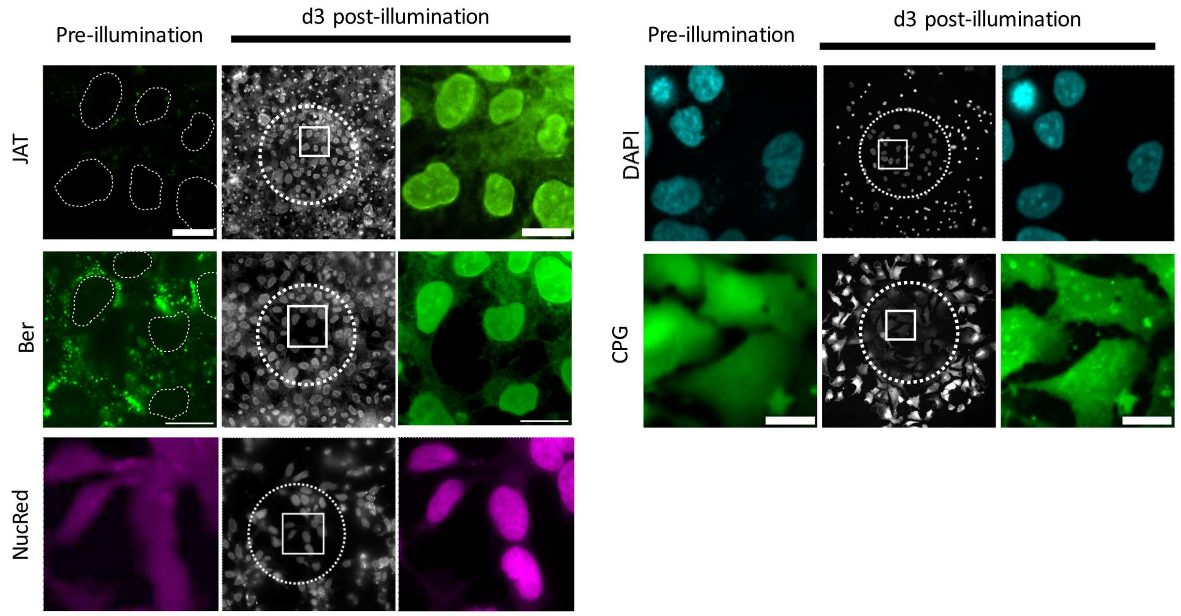


**Fig S19.** Positive optofixing mediated by nuclear fluorogenic trackers. Crops in R1 (dashed circle) is observed before irradiation (pre-irradiation) and 3 days after (d3) (scale bar: 10 µm). Results are representative of three independent experiments.

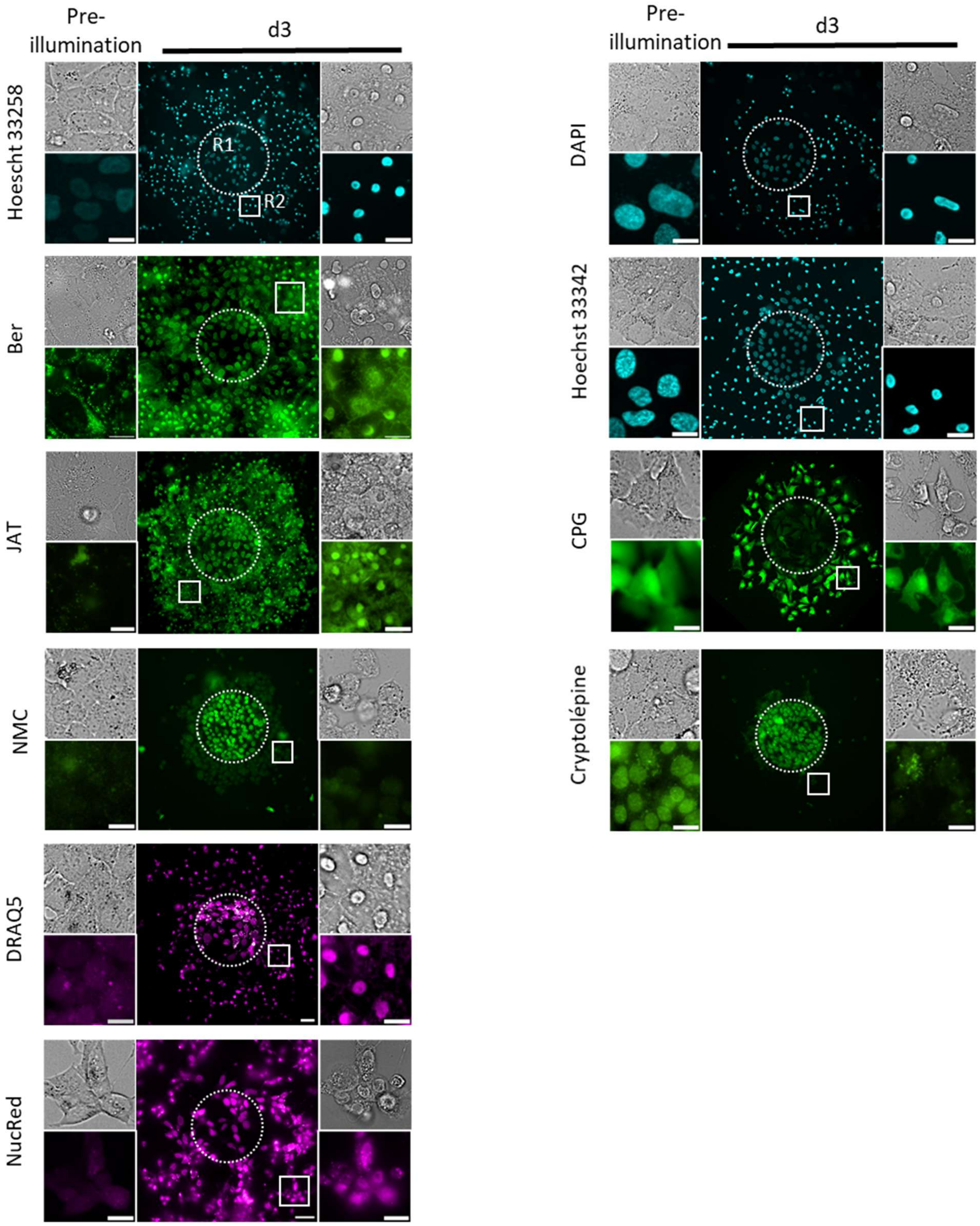


**Fig S20.** R2 region shown in optofixed cells mediated by natural or synthetic fluorophores. Optofixing reagents can be categorized as either fluorogenic (*left panel*, according to Fig. 4D) or non-fluorogenic (*right panel*, according to Fig. 4D). Crops in R2 region is observed before irradiation (pre-irradiation) and 3 days after (d3) (R1 in dashed circle) (scale bar: 10 µm). Results are representative of three independent experiments.

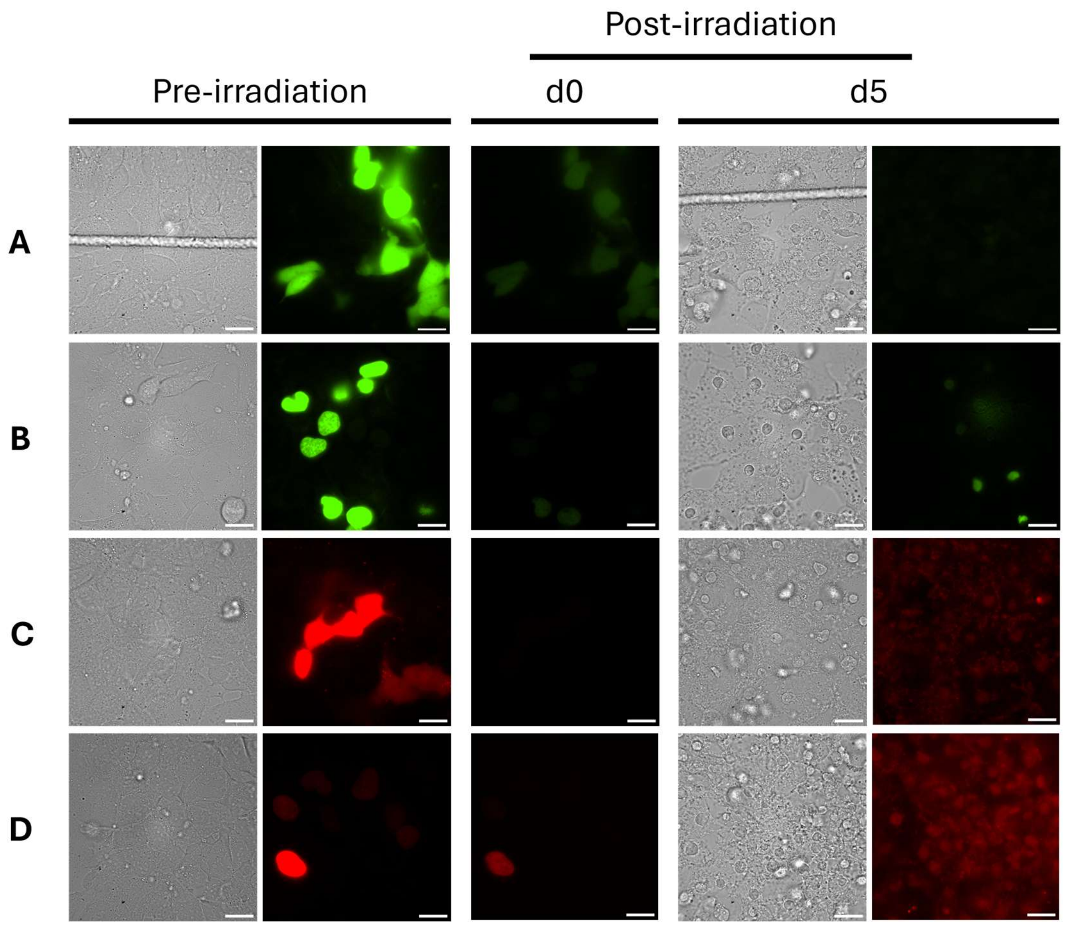


**Fig S21.** Observation of transiently transfected cells with **(A)** HA-GFP, **(B)** H2B-GFP, **(C)** HA-mCherry, **(D)** H2B-mCherry, before (pre-irradiation), immediately (d0) and 5 days after (d5) irradiation. Results are representative of two independent experiments.
